# A temporally restricted function of the Dopamine receptor Dop1R2 during memory formation

**DOI:** 10.1101/2024.05.29.594954

**Authors:** Jenifer C. Kaldun, Emanuele Calia, Ganesh Chinmai Bangalore Mukunda, Cornelia Fritsch, Nikita Komarov, Simon G. Sprecher

## Abstract

Dopamine is a crucial neuromodulator involved in many brain processes, including learning and the formation of memories. Dopamine acts through multiple receptors and controls an intricate signaling network to regulate different tasks. While the diverse functions of dopamine are intensely studied, the interplay and role of the distinct dopamine receptors to regulate different processes is less well understood. An interesting candidate is the dopamine receptor Dop1R2 (also known as Damb), as it could connect to different downstream pathways. Dop1R2 is reported to be involved in forgetting and memory maintenance, however, the circuits requiring the receptors are unknown. To study Dop1R2 and its role in specific spatial and temporal contexts, we generated a conditional knock-out line using the CRISPR-Cas9 technique. Two FRT sites were inserted, allowing flippase-mediated excision of the dopamine receptor in neurons of interest. To study the function of Dop1R2, we knocked it out conditionally in the Mushroom body of *Drosophila melanogaster*, a well-studied brain region for memory formation. We show that Dop1R2 is required for later memory forms but not for short-term aversive or appetitive memories. Moreover, Dop1R2 is specifically required in the α/β-lobe and the α’/β’-lobe but not in the γ-lobe of the Mushroom body. Our findings show a spatially and temporally restricted role of Dop1R2 in the process of memory formation highlighting the differential requirement of receptors during distinct phases of learning.

## INTRODUCTION

The neuromodulator dopamine is involved in a plethora of brain functions. Among them are learning and memory, reinforcement signaling, reward, arousal, and motor functions (Klein et al., 2019; Missale et al., 1998; Tritsch & Sabatini, 2012). Perturbations in the dopaminergic system are associated with diseases like Parkinson’s disease, addiction, depression, schizophrenia, and many more (Klein et al., 2019; Missale et al., 1998; Tritsch & Sabatini, 2012). Therefore, it is important to understand dopamine signaling in greater detail allowing to develop more effective therapeutics or preventive measures.

Dopamine is synthesized from tyrosine in dopaminergic neurons and binds to G-protein coupled receptors (GPCRs) (Klein et al., 2019; Missale et al., 1998; Yamamoto & Seto, 2014). These dopamine receptors have seven transmembrane domains, with an intracellular C-terminus and an extracellular N-terminus. The receptors interact with different G-proteins, which are heterotrimeric protein complexes consisting of an α-, β-, and γ-subunit. The G-protein complex binds to the third intracellular loop and the C-terminus of the receptors. Humans and other mammals express five different dopamine receptors, separated into two types. Type 1 dopamine receptors elevate protein kinase A (PKA) signaling and cyclic adenosine monophosphate (cAMP) levels via the alpha subunit Gα_s_, whereas type 2 receptors inhibit PKA activity via Gα_i/o_ thus reducing cAMP levels (Klein et al., 2019; Missale et al., 1998; Tritsch & Sabatini, 2012). Downstream of cAMP and PKA, transcriptional activation mediated by cAMP response element-binding protein leads to the expression of immediate early genes as a response to dopamine signaling (Carlezon et al., 2005; Neves et al., 2002). This pathway is a key regulator of long-term memory (LTM) formation (Alberini & Kandel, 2014; Kaldun & Sprecher, 2019; Kandel, 2012). Moreover, dopamine receptors can modulate internal Ca^2+^ levels by engaging the Phospholipase C signaling pathway. Furthermore, the G-protein β and γ subunits can directly modulate voltage-gated and ligand-gated ion channels (Klein et al., 2019; Ledonne & Mercuri, 2017; Missale et al., 1998; Nishi et al., 2011; Tritsch & Sabatini, 2012). However, the coordination of the receptors and the different dopamine mediated processes in specific circuits remains unexplored. Therefore, an accessible and well-characterized model is required, like the olfactory circuit of *Drosophila melanogaster*. The connectome of this circuit is known, and can be manipulated with the sophisticated genetic tools of the fly (Takemura et al., 2017; Li et al., 2020; Owald et al., 2015). Similar to mammals, the fly uses dopamine in learning, memory, forgetting, negative and positive reinforcement, locomotion, and sleep and arousal regulation (Berry et al., 2012; Burke et al., 2012; Karam et al., 2020; Sabandal et al., 2021; Sabandal et al., 2020; Siju et al., 2021; Siju et al., 2020; Sitaraman et al., 2015; Tomita et al., 2017; Waddell, 2013; Yamamoto & Seto, 2014). Additionally, L-Dopa, the precursor for dopamine and a powerful receptor agonist, as well as other pharmaceutics have been shown to function in the fly (Yamamoto & Seto, 2014). Thus, the dopaminergic system can be studied in *Drosophila*, taking advantage of the available genetic tools.

The fly uses four different dopamine receptors. Dop1R1 (Dumb) (Kim et al., 2003; Sugamori et al., 1995) and Dop1R2 (Damb) (Feng et al., 1996; Han et al., 1996) are type 1 receptors. Dop2R is a type 2 receptor (Hearn et al., 2002), and DopEcR is a type 1 dopamine receptor that also uses ecdysone as a ligand (Srivastava et al., 2005). All four receptors were shown to be involved in learning, memory and forgetting (Berry et al., 2012; Karam et al., 2020; Kim et al., 2007; Lark et al., 2017; Qi & Lee, 2014; Qin et al., 2012; Scholz-Kornehl & Schwarzel, 2016; Sun et al., 2020; Zhou et al., 2019). While it is well established that Dop1R1 is crucial for learning and short-term memory (Kim et al., 2007; Qin et al., 2012), the role of the other receptors is less clear. Dop1R2 is an interesting candidate to study, due to its ability to couple to two different G-proteins, Gα_s_, which engages the cAMP pathway, and Gα_q_, which involved in Ca^2+^-signaling (Han et al., 1996; Himmelreich et al., 2017; Sun et al., 2020). This could allow Dop1R2 to modulate learning and memory in a complex fashion. The receptor is mainly expressed in the Mushroom body (MB) (Crocker et al., 2016; Croset et al., 2018; Han et al., 1996; Kim et al., 2007; Lark et al., 2017), an important brain region for olfactory associative learning (Aso, Hattori, et al., 2014; Aso, Sitaraman, et al., 2014; Cognigni et al., 2018). Previous single cell transcriptomics data show that Dop1R2 is sparsely found in the whole nervous system (in 7.99% of ventral nerve cord cells, and in 13.8% of cells outside the MB) (Allen et al., 2020; Davie et al., 2018). Work on a mutant line suggests that Dop1R2 is involved in forgetting (Berry et al., 2012). However, a study using RNAi silencing suggests that the receptor plays a role in memory maintenance (Sun et al., 2020). As these studies used different learning assays – aversive and appetitive respectively as well as different methods, it is unclear if Dop1R2 has different functions for the different reinforcement stimuli. To resolve this problem, we generated a transgenic line to conditionally knock out *Dop1R2*, in a spatially and temporally specific manner. Using CRISPR-Cas9 and homology-directed repair (HDR), we included FRT sites in the endogenous locus of *Dop1R2* for flippase-mediated excision. In addition, we inserted an HA-Tag, to monitor the spatial localization of the receptor. This also allows us to visualize the efficiency of the flip-out. We used this line to study the role of the receptor in learning and memory in the MB for both aversive and appetitive conditioning. Upon flip-out of the receptor in the MB, 2h memory and long-term memory (LTM) are impaired. Similar results are obtained when we flip out Dop1R2 specifically in the α/β-lobes and α’/β’-lobes of the MB, which are involved in those memory phases. Therefore, Dop1R2 is required in the α/β-lobes and α’/β’-lobes for later memory forms.

## RESULTS

### Generation of the Dop1R2 conditional knock-out line

To be able to study both the requirement of Dop1R2 in specific neurons as well as the localization of the receptor we generated a transgenic line that allows defined spatial and temporal knock-out of the receptor. In short, we inserted two FRT sites for flippase-mediated excision (Golic & Lindquist, 1989; Gratz et al., 2014) as well as a 3xHA-tag to study the localization of the receptor. The HA-Tag was chosen to minimize interference with the receptor structure. A representation of the HA-tagged receptor is depicted in Figure 1A. The 3^rd^ intracellular loop between transmembrane domains (TMDs) five and six as well as the C-terminal tail are required for G-protein binding (Missale et al., 1998). Figure 1B gives an overview of the experimental strategy. The endogenous locus is cut twice by CRISPR-Cas9 to replace the first two coding exons of Dop1R2 with the corresponding sequence flanked by FRT sites and fused to the HA-Tag, using a donor plasmid as template for homolog-directed repair. The donor plasmid also contains the region directly upstream and downstream of the exchange site as homology arms to align the plasmid. The plasmid carrying the sequence for the two gRNAs as well as the donor plasmid were injected into embryos with maternal Cas9 expression (*nos>Cas9*). Using the Gal4-UAS system to express flippase in neurons of interest the receptor can be irreversibly flipped out in those neurons while keeping it functional in the rest of the brain. The first FRT site was placed in the intron before the first coding exon, which contains all seven TMDs (Figure 1C). Since *Dop1R2* has three transcript isoforms with differences in the C-terminus the second FRT site was placed at the end of the last common exon. Thus, FLP-mediated recombination will lead to deletion of the two common coding exons including all TMDs. Successful insertion was verified by sequencing.

**Fig 1.**
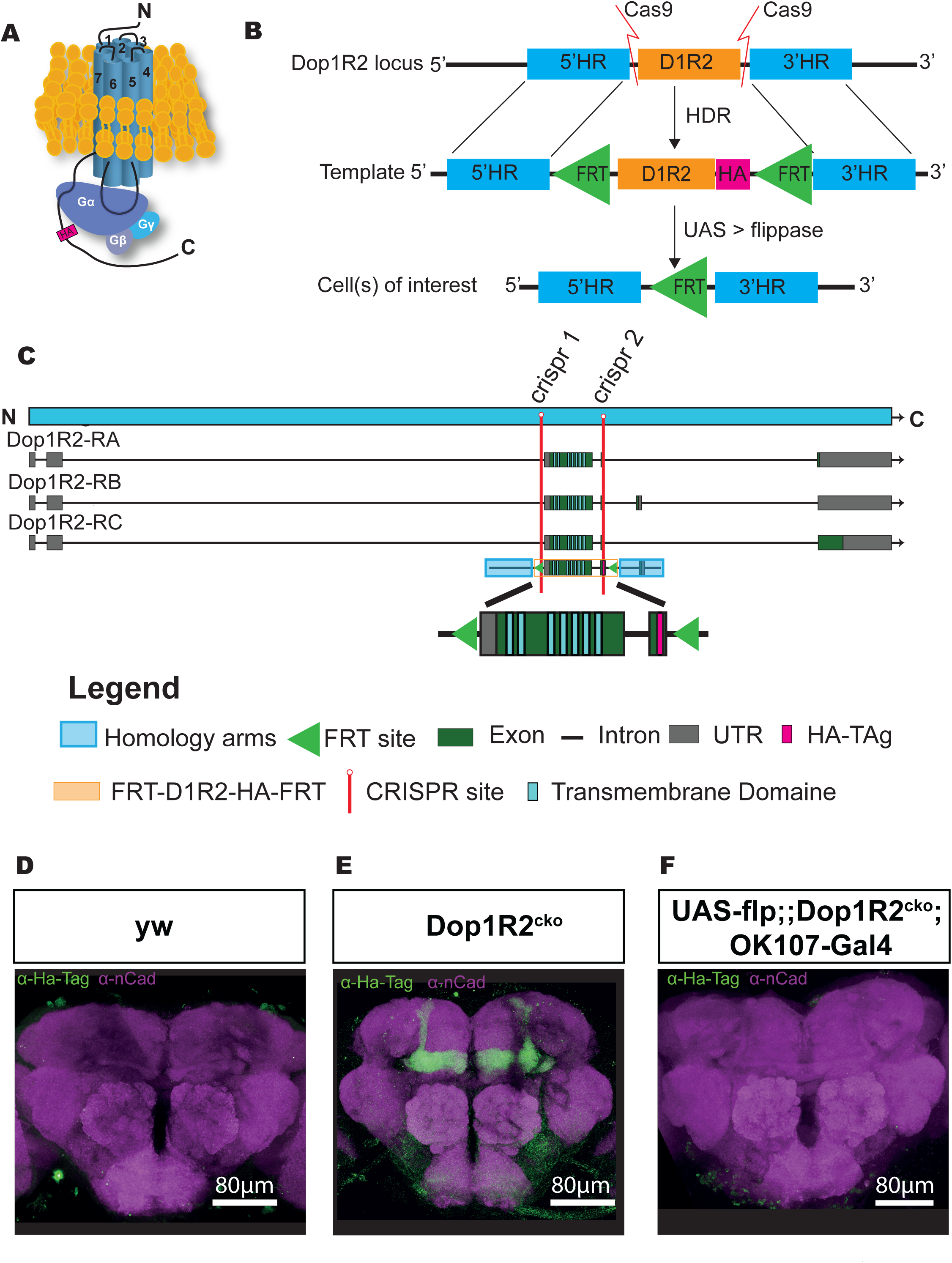
Generation of a conditional knockout allele for *Dop1R2*. A) Schematic representation of the receptor structure and interaction with the G-protein complex. The position of the HA-Tag is indicated. B) Schematic representation of the conditional knock-out system. The endogenous *Dop1R2* was replaced using CRISPR-Cas9 mediated homology-directed repair (HDR) from a donor plasmid. The plasmid contained the two common coding exons of *Dop1R2* with an HA-Tag in the C-Terminus and two FRT sites flanking this sequence. In the resulting *Dop1R2^cko^* allele the inserted HA-Tag and *Dop1R2* sequence can be removed by flippase (FLP) in cells of interest. C) Schematic representation of the *Dop1R2* gene locus with 3 different transcript isoforms. The location of the two used CRISPR sites are highlighted in red. The positions of the transmembrane domains in the isoforms and in the donor plasmid, are indicated. D-F) Dop1R2::HA expression in a frontal brain confocal section of D) y, w, E) *Dop1R2^cko^* or F) *UAS-flp/+;; dop1R2^cko^,* OK107*-*Gal4*/+* flies aged one week. The HA-Tag was visualized using an anti-HA-Tag antibody (green). Brain structures were labeled with anti-N-cadherin (nCad, magenta) antibody. Scale bar: 80 µm.

### Dop1R2^cko^ is expressed in the Mushroom body

To monitor the localization of the tagged dopamine receptors we first stained brains with an antibody against the HA-Tag in one-week-old flies. In the y, w control line (Figure 1D) the HA-tag does not show any specific staining. In the *Dop1R2^cko^* line, we see a clear signal in the entire MB (Figure 1E). This matches previous reports as well as single-cell RNAseq data showing that Dop1R2 is expressed in the MB (Croset et al., 2018; Han et al., 1996).

Flipping the receptor out in the MB using the MB-specific OK107-Gal4 driver in combination with UAS-flp abolishes the signal, demonstrating the efficiency of the FRT sites (Figure 1F). Therefore, the flip-out system seems to work as planned.

### Short-term memory is not affected by the loss of Dop1R2

We wanted to see if flipping out *Dop1R2* in the MB affects memory acquisition and STM by using classical olfactory conditioning. In short, a group of flies is presented with an odor coupled to an electric shock (aversive) or sugar (appetitive) followed by a second odor without stimulus. For assessing their memory, flies can freely choose between the odors either directly after training (STM) or at a later timepoint.

To ensure that the introduced genetic changes to the *Dop1R2* locus do not interfere with behavior we first checked the sensory responses of that line (Figure 2 S1A-D). The *Dop1R2^cko^* line shows comparable odor responses (Figure 2 S1A+B) as well as sugar and shock response (Figure 2 S1C+D) to the control line. Aversive STM (Figure 2 S1E), as well as aversive 2h memory (Figure 2 S1F), are not significantly different from the y, w control line. Moreover, appetitive STM is also comparable to the control (Figure 2 S1G). Thus, unsurprisingly, the introduced changes to *Dop1R2* do not interfere with normal receptor function.

**Fig 2.**
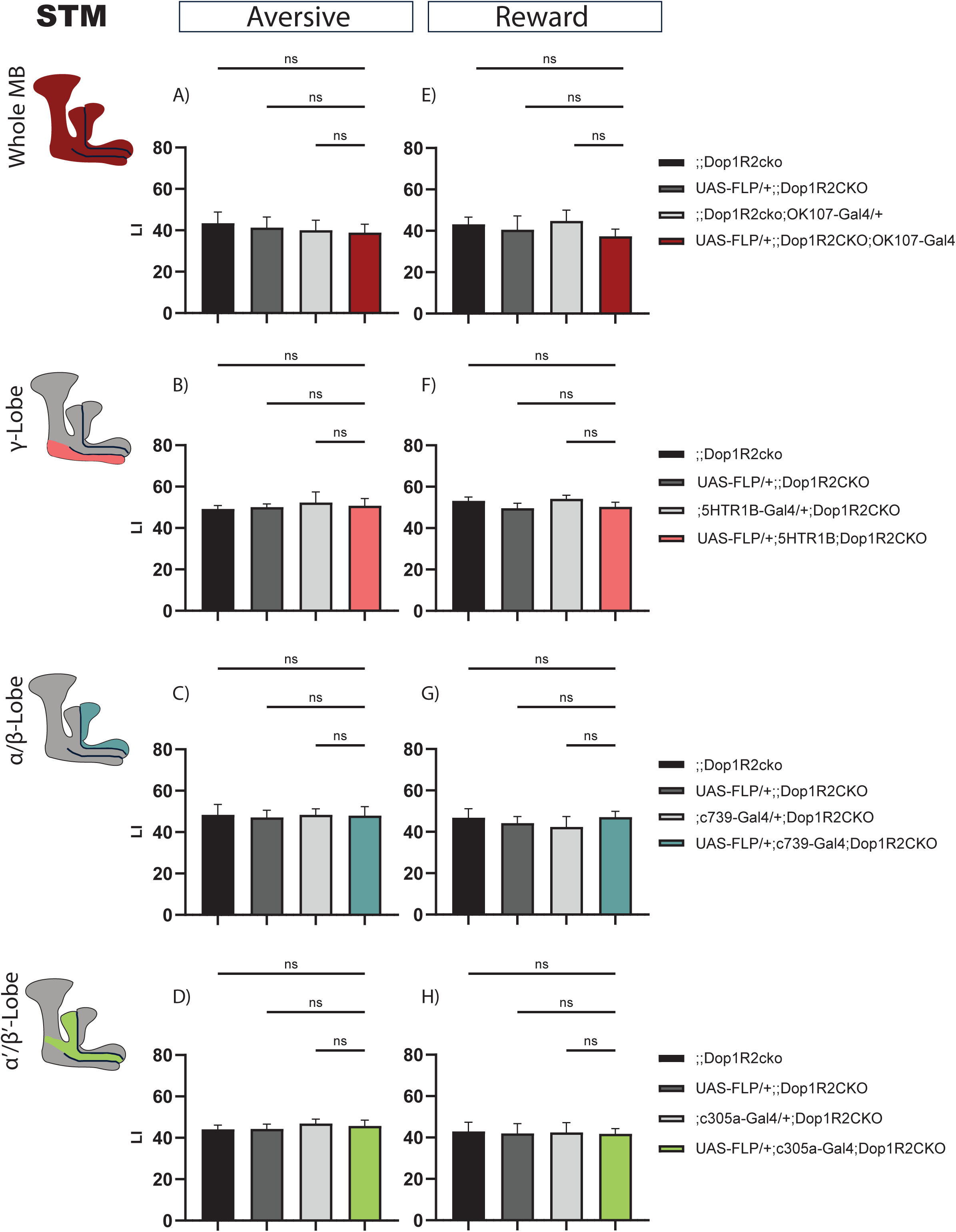
Short-term memory of flies with knock-out of Dop1R2 in the mushroom Body. A-D) Aversive training, E-H) reward training. A) and E) Whole MB flip-out using OK107*-*Gal4 and parental controls. B) and F) γ-lobe flip-out using 5HTR1B*-*Gal4 and parental controls. C) and G) α/β-lobe flip-out using c739*-*Gal4 and parental controls. D) and H) α’/β’-lobe flip-out using c305a*-*Gal4 and parental controls. No performance impairment was observed in any of the tested conditions. See Fig 2 S1 for sensory controls. Bar graphs represent the mean, and error bars represent the standard error of the mean. For each shown graph the N = 12. ns – not significant determined by a One-way ANOVA and Tukey HSD.

To test the requirement of Dop1R2 for short-term memory we assessed memory performance directly after training for flies with flipped-out *Dop1R2* in the whole MB or in individual lobes along with the parental controls. The flies were aged for a week before undergoing classical olfactory conditioning. Both aversive STM and appetitive STM were tested. But first, we ensured that the whole MB flip-out line responds normally to the used odors and stimuli. UAS-Flp/+;; Dop1R2^cko^; OK107-Gal4/+ flies show comparable responses as the parental controls to both odors (Figure 2 S1H+I), shock (Figure 2 S1J) and sugar (Figure 2 S1K).

For aversive conditioning, flipping out *Dop1R2* in the whole MB does not change STM compared to parental controls (Figure 2A). Next, we flipped out *Dop1R2* in the γ-lobe using 5HTR1B-Gal4. This MB lobe is involved in STM (Blum et al., 2009; Trannoy et al., 2011; Zars et al., 2000). Matching the results for the whole MB flip out, we do not see a change in the performance score (Figure 2B). Similar results are obtained when we flip out the receptor in the α/β-lobes using c739-Gal4 (Figure 2C) or α’/β’-lobes using c305a-Gal4 (Figure 2D).

Reward STM is also not changed when *Dop1R2* is flipped out in the whole MB (Figure 2E) in the γ-lobe (Figure 2F), the α/β-lobes (Figure 2G) or the α’/β’-lobes (Figure 2H). Taken together the results indicate that Dop1R2 is required for STM in none of the MB lobes.

### 2h memory is impaired by loss of Dop1R2

As Dop1R2 was previously described to be involved in forgetting and/or memory maintenance we wanted to assess later time points after training.

First, we looked at two hours after aversive training. Flip-out in the whole Mushroom body leads to a reduced performance score (Figure 3A). Next, we asked which MB lobe might cause this reduction. Therefore, we flipped out *Dop1R2* in the γ-lobe using the 5HTR1B-Gal4 driver, characterized in Aso et al., 2012, in the α/β-lobes using c739-Gal4, or α’/β’-lobes using c305a-Gal4. Loss of Dop1R2 in the γ-lobe does not reduce the memory performance (Figure 3B). However, loss of Dop1R2 in both the α/β-lobes (Figure 3C) or the α’/β’-lobes (Figure 3D) impaired 2h memory after aversive training. We tested the same for reward memory using sugar as reinforcement. Flip out of *Dop1R2* in the whole MB (Figure 3E) results in a reduced performance score. Loss of Dop1R2 in the γ-lobe does not affect memory performance (Figure 3F). As for aversive training, flip out of *Dop1R2* in the α/β-lobes (Figure 3G) or the α’/β’-lobes (Figure 3H) impaired 2h memory after reward training. However, the reduction is not as severe as for aversive training. Taken together the results indicate that Dop1R2 is required for aversive 2h memory as well as reward 2h memory in the α/β-lobes and the α’/β’-lobes but is dispensable in the γ-lobe.

**Fig 3.**
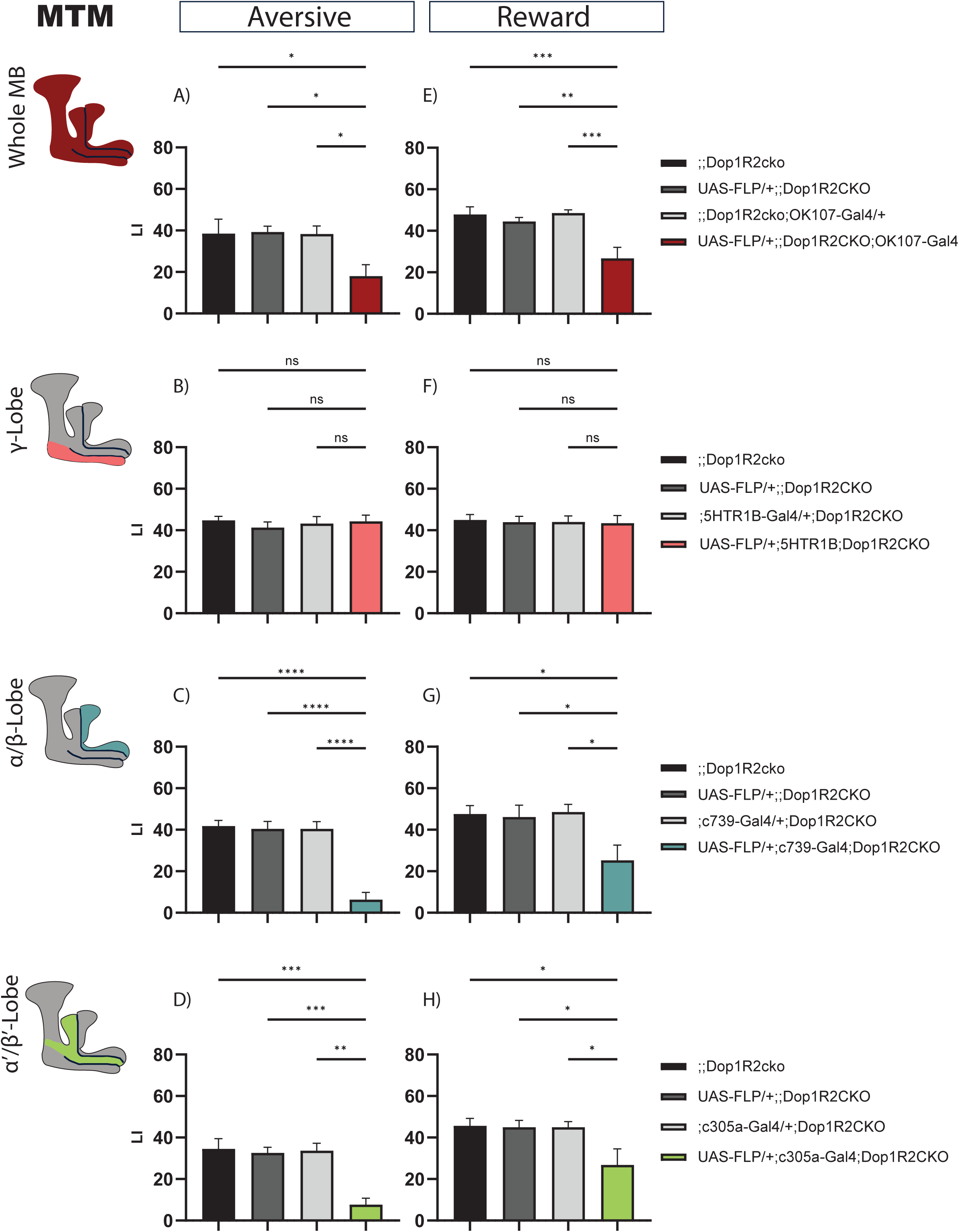
2h memory of flies with knock-out of Dop1R2 in the mushroom Body. A-D) Aversive training, E-H) reward training. A) and E) Whole MB flip-out using OK107*-*Gal4 and parental controls. B) and F) γ-lobe flip-out using 5HTR1B*-*Gal4 and parental controls. C) and G) α/β-lobe flip-out using c739*-*Gal4 and parental controls. D) and H) α’/β’-lobe flip-out using c305a*-*Gal4 and parental controls. For whole MB flip-out, α/β-lobes and α’/β’-lobes both aversive and appetitive 2h memory performance is impaired. Loss of Dop1R2 in the γ-lobe does not affect 2h memory. See Fig 2 S1 for sensory controls. Bar graphs represent the mean, and error bars represent the standard error of the mean. For each shown graph the N = 12. Asterisks denote significant differences between groups (*p < 0.05, **p < 0.005, ***p < 0.001, ****p < 0.0001, ns: not significant) determined by One way ANOVA and Tukey HSD (Panels A-C, E-H) and Kruskal-Wallis with Dunn’s multiple comparisons test due to non-normal distribution (Panel D).

### 24h memory is impaired by loss of Dop1R2

Next, we wanted to see if later memory forms are also affected. One cycle of reward training is sufficient to create LTM (Krashes & Waddell, 2008), while for aversive memory, 5-6 cycles of electroshock-trainings are required to obtain robust long-term memory scores (Tully et al., 1994). So, we looked at both 24h aversive and appetitive memory. For aversive LTM, the flies were tested on the Y-Maze apparatus as described in Mohandasan et al., 2022.

Flipping out *Dop1R2* in the whole MB causes a reduced 24h memory performance (Figure 4A, E). No phenotype was observed when *Dop1R2* was flipped out in the γ-lobe (Figure 4B, F). However, similar to 2h memory, loss of Dop1R2 in the α/β-lobes (Figure 4C, G) or the α’/β’-lobes (Figure 4D, H) causes a reduction in memory performance. Thus, Dop1R2 seems to be involved in aversive and appetitive LTM in the α/β-lobes and the α’/β’-lobes.

**Fig 4.**
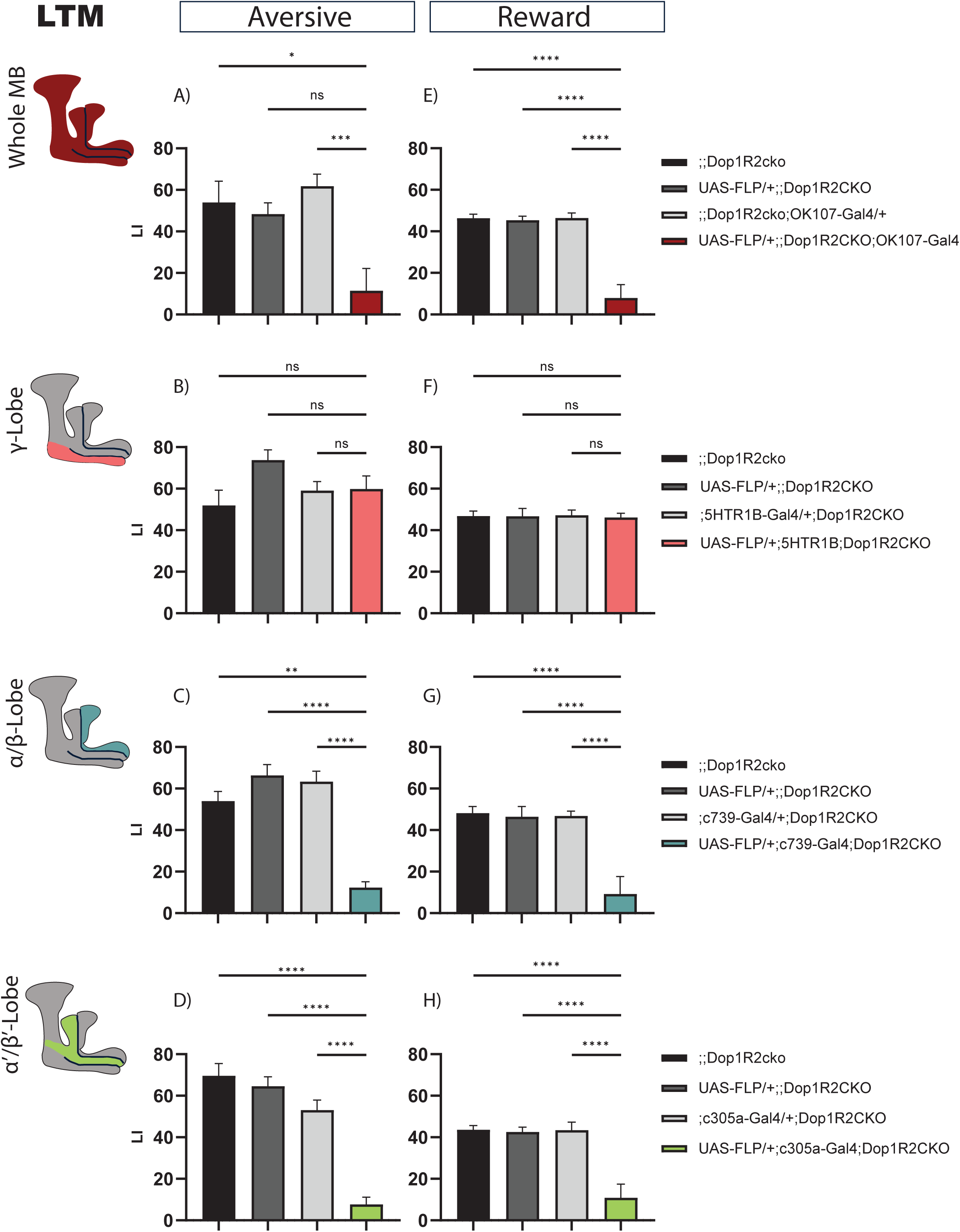
24h memory of flies with knock-out of Dop1R2 in the mushroom Body. A-D) Aversive training, E-H) reward training. A) and E) Whole MB flip-out using OK107*-* Gal4 and parental controls. B) and F) γ-lobe flip-out using 5HTR1B*-*Gal4 and parental controls. C) and G) α/β-lobe flip-out using c739*-*Gal4 and parental controls. D) and H) α’/β’-lobe flip-out using c305a*-*Gal4 and parental controls. For whole MB flip-out, α/β-lobes and α’/β’-lobes aversive and appetitive 24h memory performance is impaired. Loss of Dop1R2 in the γ-lobe does not affect 24h memory. See Fig 2 S1 for sensory controls. Bar graphs represent the mean, and error bars represent the standard error of the mean. For each shown graph included in the reward training experiment the N = 12, while for the graphs included in the aversive training experiment the N = 14. Asterisks denote significant differences between groups (*p < 0.05, **p < 0.005, ***p < 0.001, ****p < 0.0001, ns: not significant determined by Kruskal-Wallis with Dunn’s multiple comparisons test due to non-normal distribution (Panels A-C) and One way ANOVA and Tukey HSD (Panels D-H).

Previous studies have shown mutation in the Dop1R2 receptor leads to improvement in LTM when a single shock training paradigm is used (Berry et al., 2012). As we found that it disrupts LTM, we wanted to verify if the absence of Dop1R2 outside the MB is what leads to an improvement in memory. To that extent, we tested pan-neuronal flip-out of Dop1R2 flies for 6hr and 24hr memory upon single shock using the elav-Gal4 driver. We found that it did not improve memory at both time points (Figure 4 S1). Confirming that flipping out Dop1R2 pan-neuronally does not improve LTM (Figure 4 S1C) and highlighting its irrelevance in memory outside the MB.

### Developmental defects are ruled out in a temporally restricted Dop1R2 conditional knockout

To exclude developmental defects in the MB caused by flip-out of Dop1R2, we stained fly brains with a FasII antibody. Compared to genetic controls, the anatomical organization as judged by FasII staining in flies lacking Dop1R2 in the mushroom body was not altered (Figure 4 S2C).

To further provide behavioral evidence that the learning defects we observed are not due to developmental knock-out of Dop1R2, we generated a Gal80^ts^-containing line, enabling the temporal control of Dop1R2 knockout in the entire mushroom body (MB). Given that the half-life of the receptor remains unknown, we assessed both aversive short-term memory (STM) and long-term memory (LTM) to determine whether post-eclosion ablation of Dop1R2 in the MB produced differences compared to our previously tested line, in which Dop1R2 was constitutively knocked out from fertilization. To achieve this, flies were maintained at 18°C until eclosion and subsequently shifted to 30°C for five to seven days. On the fifth day, training was conducted, followed by memory testing. Our results indicate that aversive STM was not significantly impaired in Dop1R2-deficient MBs compared to control flies (Figure 4 S3), consistent with our previous findings (Figure 2). However, aversive LTM was significantly impaired relative to control lines (Figure 4 S3), which also aligned with prior observations. These findings strongly indicate that memory loss caused by Dop1R2 flip-out is not due to developmental defects.

## Discussion

We have generated a conditional knock-out line for the dopamine receptor Dop1R2 following a similar approach as in Widmer et al., 2018. To achieve this, FRT sites were inserted in front of the start codon and in the C-Terminus of Dop1R2 using CRISPR-Cas9 mediated homology-directed repair. In addition, an HA-Tag was inserted to be able to visualize the dopamine receptor expression as well as verify successful flip out. Using an anti-HA-Tag antibody we were able to visualize Dop1R2 in the MB of the generated line. This matches previous reports for the expression of *Dop1R2* (Crocker et al., 2016; Croset et al., 2018; Han et al., 1996; Kim et al., 2007; Sun et al., 2020). Moreover, upon knocking out *Dop1R2* with an MB-specific driver, the HA-tag labeling disappears indicating that the conditional knock-out system works. The HA-Tag could also be useful to study the subcellular localization of Dop1R2 within the MB lobes, for example if it is close to synapses of dopaminergic neurons.

To get a better overview of Dop1R2’s role in the Mushroom body we analyzed appetitive memory at different timepoints after training in the individual MB lobes. Loss of Dop1R2 in the whole MB as well as the α/β-lobe or the α’/β’-lobe impairs 24h reward memory. This observation matches previous studies. Using *Dop1R2-RNAi* in the MB, Sun et al., 2020 showed that STM is intact, while LTM is impaired. When knocking down the Raf/MAPK pathway they get a similar phenotype. Moreover, expression of a constitutive active *Raf* allele rescues the Dop1R2 dependent LTM deficit and Dop1R2 seems to be required for the phosphorylation of the MAPK. Therefore, Dop1R2 dependent activation of the Raf/MAPK pathway is required for stabilization of reward LTM memory. Another study shows that reward LTM requires food with a high energetic value (Musso et al., 2015). This signal seems to be relayed by the dopaminergic neuron (DAN) MB-MP1 (PPL1-γ1pedc), that showed temporally restricted oscillating activity early post-training (Musso et al., 2015; Pavlowsky et al., 2018). Furthermore, knock-down of Dop1R2 using RNAi impaired reward LTM while leaving STM intact. Therefore, dopaminergic signaling through MB-MP-1 and Dop1R2 could indicate the energetic value of the reward and decide if LTM should be formed or not. Interestingly, the MB-MP1 DAN arborizes in the spur of the γ-lobe as well as the inner core of the pedunculus, which consists of the axons of the α/β Kenyon Cells (KCs) (Tanaka et al., 2008), which according to our results require Dop1R2 for reward LTM. Loss of Dop1R2 in the Mushroom Body Output neuron MVP2 (MBON-γ1pedc>α/β) also impaired reward LTM while leaving STM intact (Pavlowsky et al., 2018). This GABAergic MVP2 MBON forms a feedback circuit with the MB-MP1 DAN. After training, the oscillatory activity of MB-MP1 is enhanced, while MVP2 is inhibited. After 30min, MVP2 gets activated and MB-MP1 is inhibited. Moreover, Dop1R2 seems to be required for modulating this feedback loop.

The MBONs are an important contributor to memory. Moreover, MBONs receive dopaminergic input and seem to express dopamine receptors (Crocker et al., 2016). We showed that Dop1R2 is required in the α/β-lobe and the α’/β’-lobe for reward LTM formation. Interestingly, MBONs which are involved in reward LTM (Ichinose et al., 2015; Owald et al., 2015; Plaçais et al., 2013) have arborization in the α/β-lobe and the α’/β’-lobe. Thus, Dop1R2 could modulate these connections as well.

In all, Dop1R2 is required for reward LTM formation and loss of the receptor impairs LTM. Dop1R2 seems to influence reward LTM in different ways. Firstly, by acting on the Raf-MAPK pathway to stabilize memory, secondly by relaying the energetic content of the reward and third, by modulating the MB-MP1-MVP2 loop.

For aversive conditioning, we observe that loss of Dop1R2 in the MB leads to impaired 2h memory and LTM, whereas STM is intact. Moreover, Dop1R2 seems to be required in the α/β-lobe and the α’/β’-lobe. A previous study using a mutant for *Dop1R2*, that also affects the neighboring gene *GC1907*, observed higher memory retention (Berry et al., 2012). Further, lack of Dop1R2 impairs reversal learning. It is proposed that Dop1R2 acts on the RAC-forgetting pathway (Cervantes-Sandoval et al., 2016; Shuai et al., 2010). Interestingly, Dop1R2 can act on two different downstream second messenger systems by using two different G-proteins (Himmelreich et al., 2017). By coupling to G_αs_ the cAMP pathway is activated. By coupling to G_αq_ the Ca^2+^ messenger system is activated. Furthermore, knocking down G_αq_ pan-neuronally or in the MB leads to memory enhancement 3h after training, but not 6h after training (Himmelreich et al., 2017). Thus, Dop1R2 could regulate the RAC forgetting pathway through G_αq_. Some RNA binding proteins and Immediate early genes help maintain identities of Mushroom body cells and are regulators of local transcription and translation (de Queiroz et al., 2025; Raun et al., 2025). So, the availability of different G-proteins may change in different lobes and during different phases of memory. The G-protein via which GPCRs signal may depend on the pool of available G-proteins in the cell/sub-cellular region (Hermans, 2003). Therefore, Dop1R2 may signal via different G-proteins in different compartments of the Mushroom body and also different compartments of the neuron. We looked at G_αo_ and G_αq_ as they are known to have roles in learning and forgetting (Ferris et al., 2006; Himmelreich et al., 2017). We found that Dop1R2 co-expresses more frequently with G_αo_ than with G_αq_ (Figure 1 S1). While there is evidence for Dop1R2 to act via G_αq_ (Himmelreich et al., 2017). It is difficult to determine whether this interaction is exclusive, or if Dop1R2 can also be coupled to other G-proteins. It will be interesting to determine the breadth of G-proteins that are involved in Dop1R2 signaling.

Berry and colleagues also observed that blocking output of DANs inhibits forgetting, while activating DANs accelerates forgetting (Berry et al., 2012). This modulation seems to require ongoing activity of the MP1 DAN together with further DANs. Interestingly, using a spaced training protocol and a *Dop1R2* RNAi knockdown, another study showed impaired LTM (Placais et al., 2017). Furthermore, after spaced training, flies have a higher energy uptake and the energy metabolism is upregulated in the MB. This increase in energy consumption is mediated by dopaminergic signaling from the MB-MP1 DAN. Loss of Dop1R2 in the MB abolishes the increase in energy consumption. Therefore, Dop1R2 seems to be important for aversive LTM formation by regulating energy consumption. Thus, like for reward LTM formation, aversive LTM seems to require sufficient energy. Starved flies reduce the formation of aversive LTM (Plaçais & Preat, 2013; Plaçais et al., 2012). This information seems to be relayed through ongoing oscillation of MB-MP1 and Dop1R2 after training. In addition, the MVP2 MBON might also be involved (Ueoka et al., 2017). Therefore, the gating mechanism in both aversive and reward LTM formation seems to require Dop1R2 (Pavlowsky et al., 2018).

In addition, the MAPK signaling pathway might also be required in aversive LTM formation by activating transcription factors like CREB and c-fos (Miyashita et al., 2018).

Both Dop1R2 and the ongoing activity of MB-MP1 seem to have multiple roles directly after training in a short time window (Berry et al., 2012; Placais et al., 2017; Plaçais et al., 2012). The circuit acts like a gating mechanism for LTM to ensure, that there is enough energy for continuing LTM formation. The MBON MVP2 acts as feedback loop to regulate the activity of the DAN. Moreover, both the RAC and the Raf/MAPK signaling pathway seem to be engaged to either forget or stabilize the memory. In aversive memory formation, loss of Dop1R2 could lead to enhanced or impaired memory, depending on the activated signaling pathways. The signaling pathway that is activated further depends on the available pool of secondary messengers in the cell (Hermans, 2003) which may be regulated by the internal state of the animal. However, it remains unclear, how all of these aspects are integrated and if there is a hierarchical order.

DANs can produce aversive and appetitive associations depending on the temporal presentation of odor cue and reinforcement stimulus (Handler et al., 2019). Thus, dopaminergic signaling can modify the KC-MBON synapses bi-directionally. The Dop1R1-G_αs_-cAMP pathway seems to detect the temporal coincidence of the stimuli whereas the Dop1R2-G_αq_-Ca^2+^ pathway detects the temporal ordering (Handler et al., 2019). Dop1R2 mutant flies seem to be able to form an odor association but are not able to update it. Further, Dop1R2 is required for the potentiation of the KC-MBON synapse following backward pairing.

Taken together, Dop1R2 has multiple roles during memory formation and integrates different signals, including the detection of the order of stimuli, the internal state, like energy levels and forgetting and maintenance signals.

The receptor does not seem to be required for STM but for later timepoints. Previous studies looking at the temporal requirement of the lobes (Guven-Ozkan & Davis, 2014; Perisse et al., 2013) defined the γ-lobe to be responsible for memory acquisition (Blum et al., 2009; Trannoy et al., 2011; Zars et al., 2000). The α/β-lobe and its output is involved in LTM (Akalal et al., 2011; Blum et al., 2009; Cervantes-Sandoval et al., 2013; Huang et al., 2012; Ichinose et al., 2015; Krashes & Waddell, 2008; Trannoy et al., 2011). The function of the α’/β’-lobe seems to be LTM consolidation (Cervantes-Sandoval et al., 2013; Krashes & Waddell, 2008). However, both lobes also seem to have a role in middle-term memory (Bouzaiane et al., 2015; Scheunemann et al., 2012; Shyu et al., 2019; Turrel et al., 2022). As loss of Dop1R2 in the γ-lobe or the whole MB does not impair STM, we conclude that the receptor is not required for this memory type. However, *Dop1R2* is expressed in the γ-lobe (Crocker et al., 2016; Croset et al., 2018), so it might be required for other types of behaviors.

The impairment of reward LTM upon knock-out of *Dop1R2* in the α/β-lobe and the α’/β’-lobe matches the described role of these neurons. So, both the α/β-lobe and the α’/β’-lobe require Dop1R2 for LTM. Interestingly, both lobes also seem to require Dop1R2 for two-hour memories. As the MB-MP1 DAN is active in this time window as well, this would suggest that Dop1R2 function at this time point is important for correct LTM formation.

This would indicate, that Dop1R1 is the main contributor to STM while Dop1R2 is responsible for later memory stages. It would be interesting to know how the switch from Dop1R1 dependency to Dop1R2 occurs.

As Dop1R2 is required in the MBON MVP2 for reward LTM, it would be exciting to see if it is also required in other MBONs or neurons outside the MB. Besides learning and memory, the MB also uses dopamine to regulate sleep. This tool offers the opportunity to study both aspects in neurons of interest.

The genetic tool we generated here to study the role of the Dop1R2 dopamine receptor in cells of interest, is not only a good substitute for RNAi knockouts, which are known to be less efficient in insects and in many instances may cause a partial, hypomorphic phenotype (Joga et al., 2016), but also provides versatile possibilities as it can be used in combination with the powerful genetic tools of *Drosophila*.

Using this line, we could show, that Dop1R2 is specifically required for later memory stages of both aversive and appetitive memory in the α/β-lobe and the α’/β’-lobe.

## Acknowledgments

We would like to thank H. Tanimoto, the Kyoto and Bloomington stock centers for fly strains. We would like to thank Dr Christine Guzman for help with transcriptomics analyses, Christopher Aeschbacher and the Bioimage Core Facility for help with imaging. We would like to thank colleagues of the Sprecher lab for their valuable input and discussions.

## Author Contribution

Conceptualization: SGS, Data curation: JCK, GCBM, EC, Formal Analysis: JCK, GCBM, EC, Funding acquisition: SGS, Investigation: JCK, NK, CF, GCBM, EC, Methodology: JCK, CF, GCBM, EC, Project administration: SGS, Resources: SGS, Supervision: SGS, Validation, SGS, CF, Visualization: JCK, NK, GCBM, EC Writing – original draft: JCK, Writing – revised draft – JCK, NK, CF, GCBM, EC, SGS.

## Supplementary FIGURE LEGENDS

**Figure 1 Supplementary 1.**
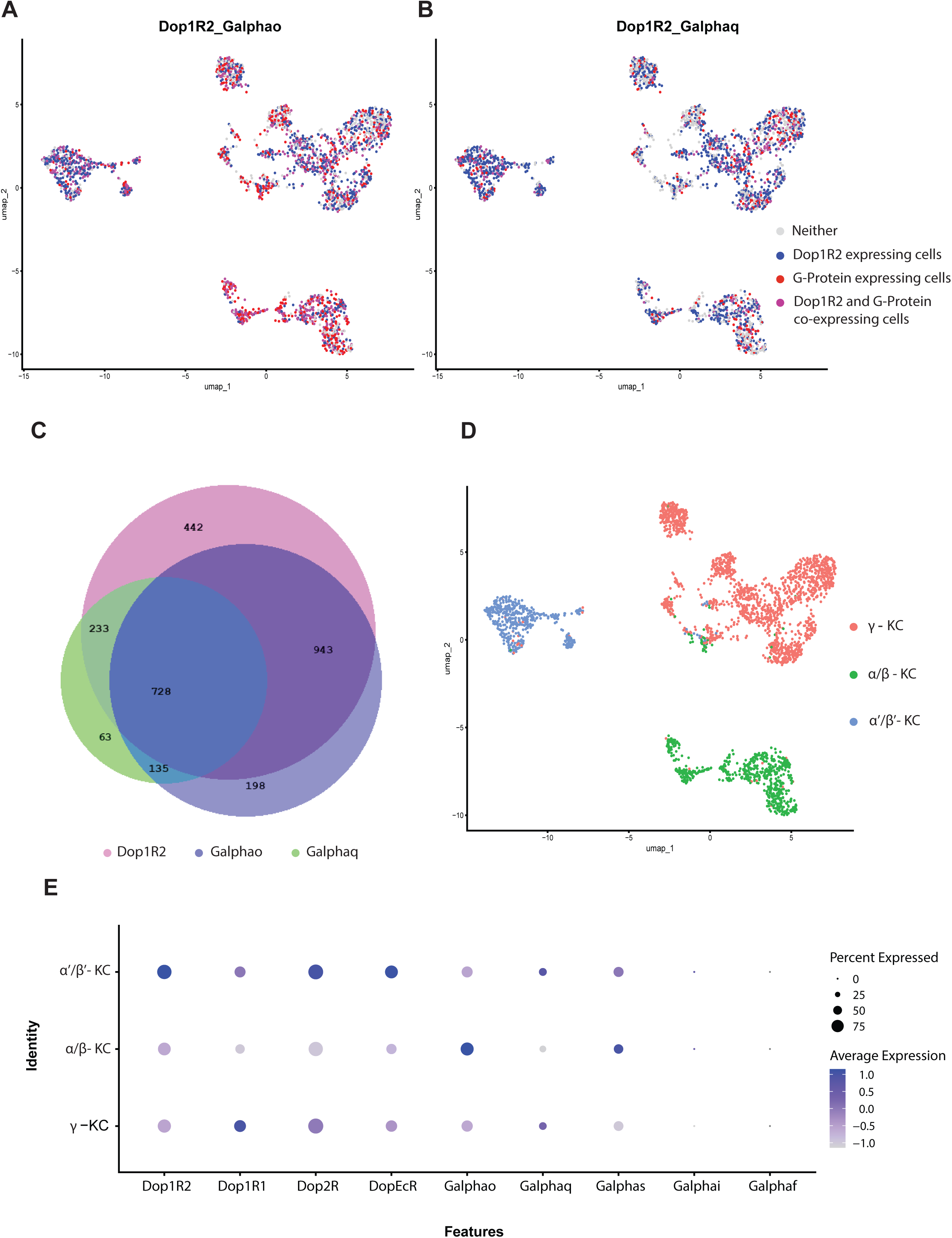
Single-cell Transcriptomic analysis of Dop1R2 and various G-proteins in the Mushroom Body neurons. A-B) Co-expression of Dop1R2 and G_αo_ and G_αq_ respectively. Cells expressing Dop1R2 are depicted in red, while the G-proteins are depicted in blue. While cells expressing both Dop1R2 and one of the G-proteins is shown in magenta. C) Venn Diagram of cells expressing Dop1R2, G_αo_ and G_αq_ in pink, violent and green respectively. The size of the circles corresponds to the number of cells expressing proteins. D) umap clustering of Mushroom body neurons, γ-lobe neurons are shown in pink, α/β-lobe neurons are shown in green while α’/β’-lobe neurons are blue. E) DotPlot of Dopamine receptors and G-Proteins in the different MB lobes. The color and size of the dots represent the average expression and percentage of expression respectively.

**Figure 2 Supplement 1.**
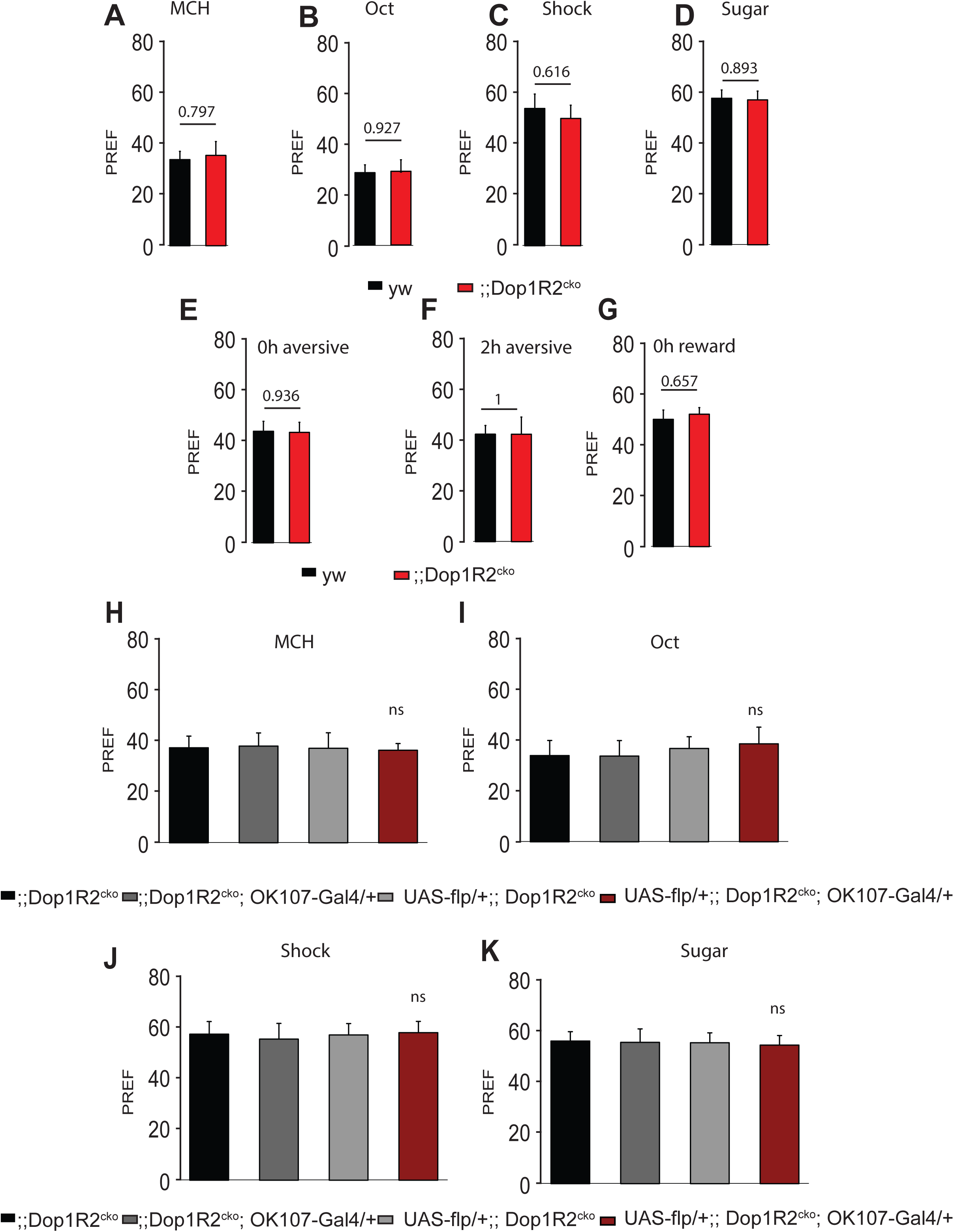
Sensory tests of the *Dop1R2* conditional knock-out line. A) Response of *y w* and *Dop1R2^cko^* to MCH. B) Response of *y w* and *Dop1R2^cko^* to Oct. A) Response of *y w* and *Dop1R2^cko^* to Shock. A) Response of *y w* and *Dop1R2^cko^* to Sugar. E) 0h aversive memory of *y w* and *Dop1R2^cko^*. F) 2h aversive memory of *y w* and *Dop1R2^cko^*. G) 0h reward memory of *y w* and *Dop1R2^cko^*. In all tested conditions the *Dop1R2^cko^* shows no significant difference to the control line. H-K) Sensory responses of flies with *Dop1R2* knock-out in the whole MB alongside the parental control lines. H) Response to MCH, I) Response to Oct, J) Response to Shock, K) Response to Sugar. Loss of Dop1R2 in the whole MB does not affect sensory responses. See S1 Table for the data. Bar graphs represent the mean, and error bars represent the standard error of the mean. For each shown graph the N = 12. Asterisks denote significant differences between groups (*p < 0.05, **p < 0.005, ***p < 0.001, ns: not significant) determined by One way ANOVA and Tukey HSD.

**Figure 4 Supplement 1.**
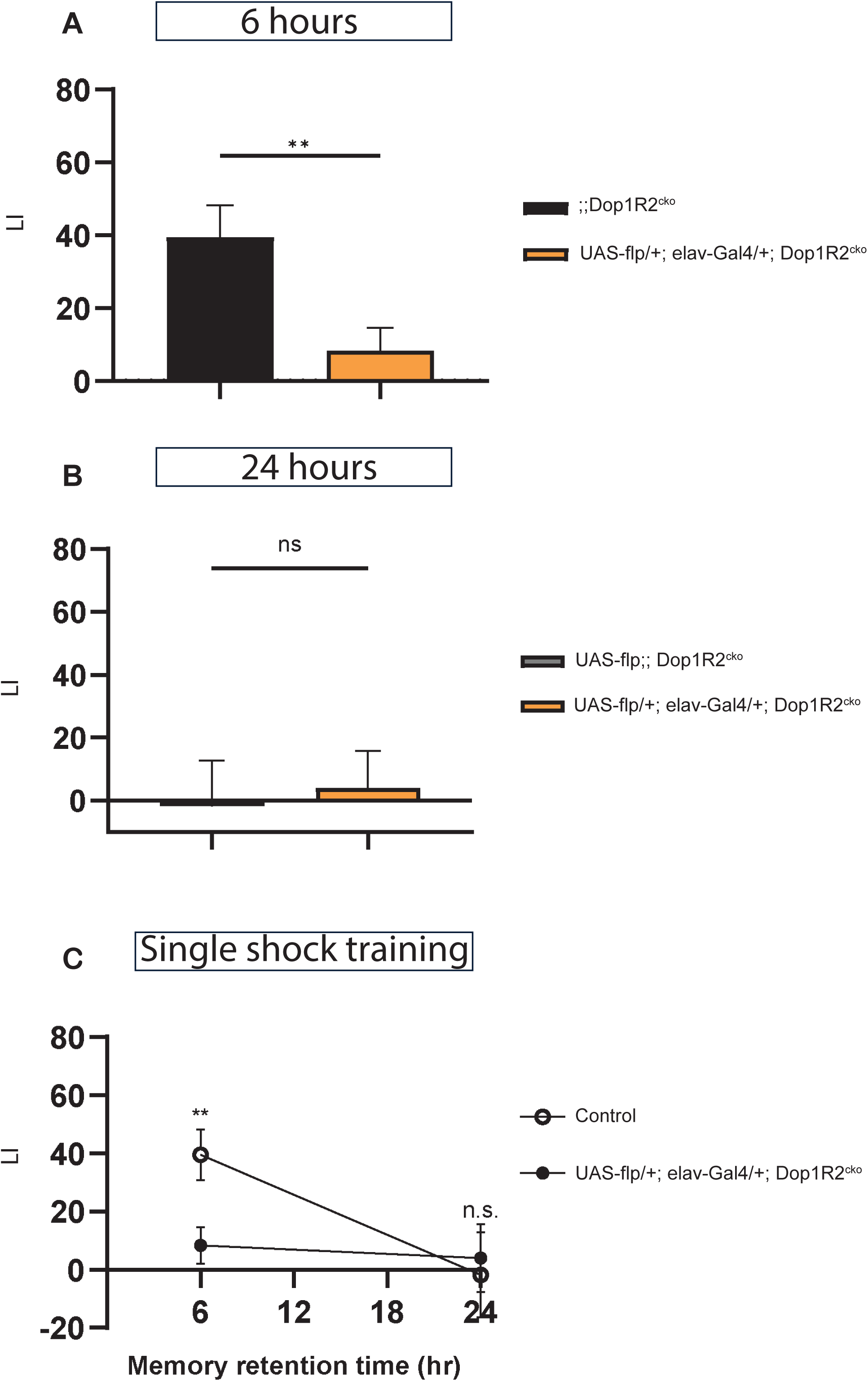
Memory retention upon single shock aversive training in *Dop1R2* pan-neuronal knockout flies. A) 6 hours memory performance of *Dop1R2* pan-neuronal knockout flies using elav*-*Gal4. B) 24 hours memory performance of *Dop1R2* pan-neuronal knockout flies using elav*-*Gal4. C) Memory retention over 24 hours in *Dop1R2* pan-neuronal knockout flies. Lack of the receptor in the whole nervous system does not improve memory retention suggesting that *Dop1R2* sparsely expressed outside of the MB might not be regulating any memory process. Single shock training is not sufficient to retain aversive memory for 24 hours in control flies either. For each graph shown the N = 14. Asterisks denote significant differences between groups (*p < 0.05, **p < 0.005, ***p < 0.001, ns: p>0.05) determined by unpaired t-test. The one-sample t-test showed no significant difference from zero for all the groups apart from the control line in the 6 hours memory experiment.

**Figure 4 Supplement 2.**
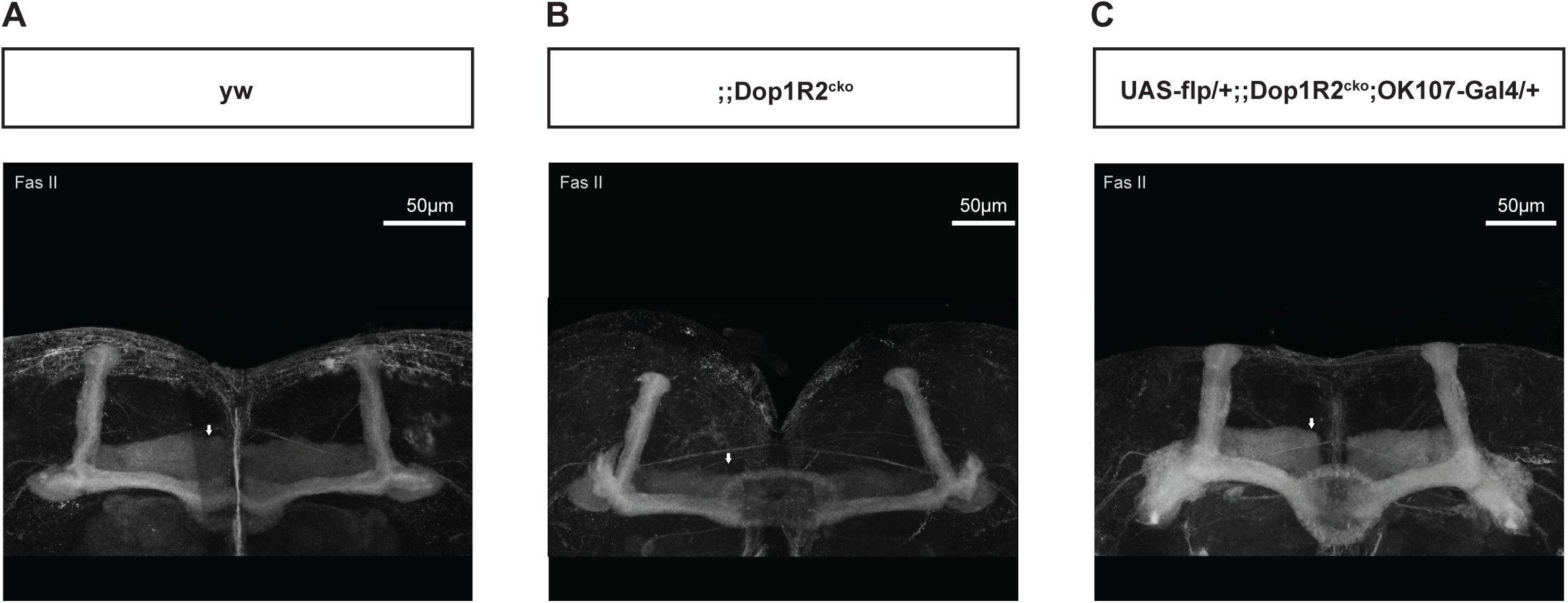
Gamma lobe development in the *Dop1R2^cko^* carrying flies. A-C) FasII expression in a frontal view of whole brain immunostaining, highlighting the Mushroom body. The *Dop1R2^cko^* modification does not affect gamma lobe (white arrows) development. Images were taken at 63x magnification, Scale bar: 50 µm.

**Figure 4 Supplement 3.**
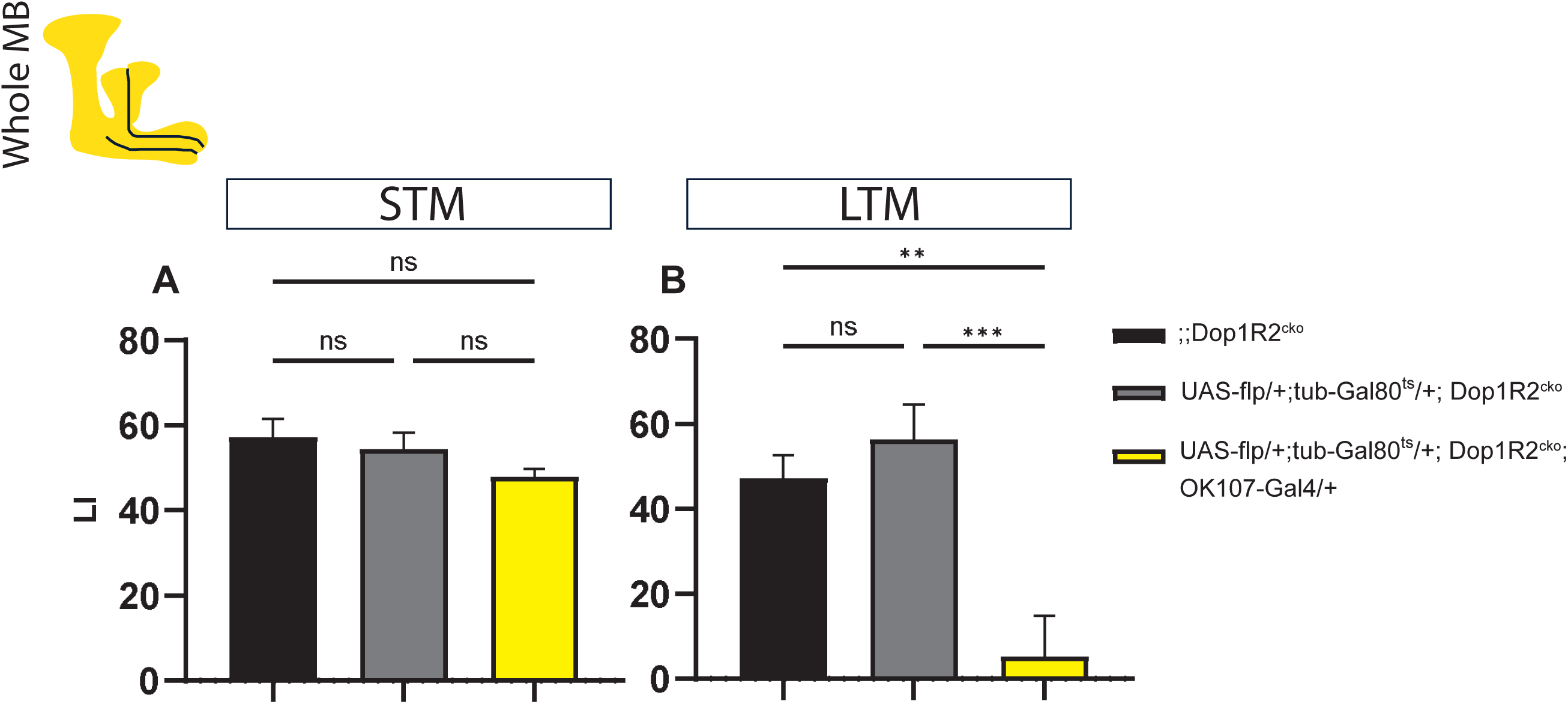
A temporally restricted knock-out of Dop1R2 in the mushroom Body rules out developmental defects. A) and B) Aversive training of flies with whole Mushroom Body flip-out at 0 days post-eclosion, using the flies ;;Dop1R2^cko^ and UAS-flp/+; tub-Gal80^ts^/+; Dop1R2^cko^ as control and UAS-flp/+;tub-Gal80^ts^/+;Dop1R2^cko^; OK107-Gal4/+ the experimental, temporally restricted knockout flies. A) Short term memory tested 0 hours after training. B) Long term memory tested 24 hours after training. Loss of Dop1R2 in the whole mushroom body exclusively after eclosion had no effect on STM but significantly impaired memory retention at 24 hours. Bar graphs represent the mean, error bars represent the standard error of the mean. For the STM graph the N = 9, while for the LTM graph the N = 14. Asterisks denote significant differences between groups (*p < 0.05, **p < 0.005, ***p < 0.001, ****p < 0.0001, ns: not significant) determined by One way ANOVA and Tukey HSD

## METHODS

### Lead Contact and Data Availability

Further information and requests for resources and reagents should be directed to and will be fulfilled by the Lead Contact, Simon Sprecher. All used data is in this manuscript.

**Table 1:**
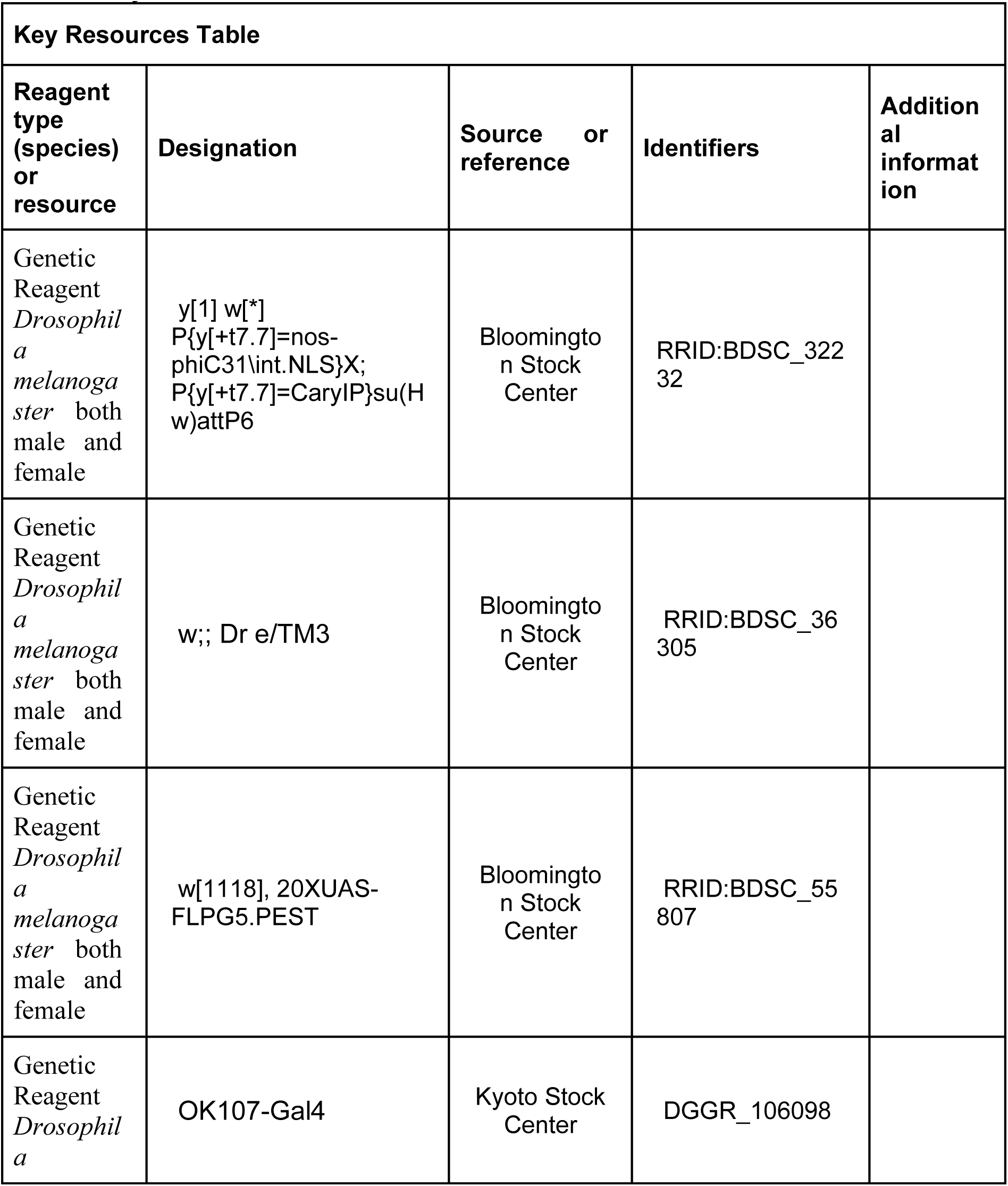
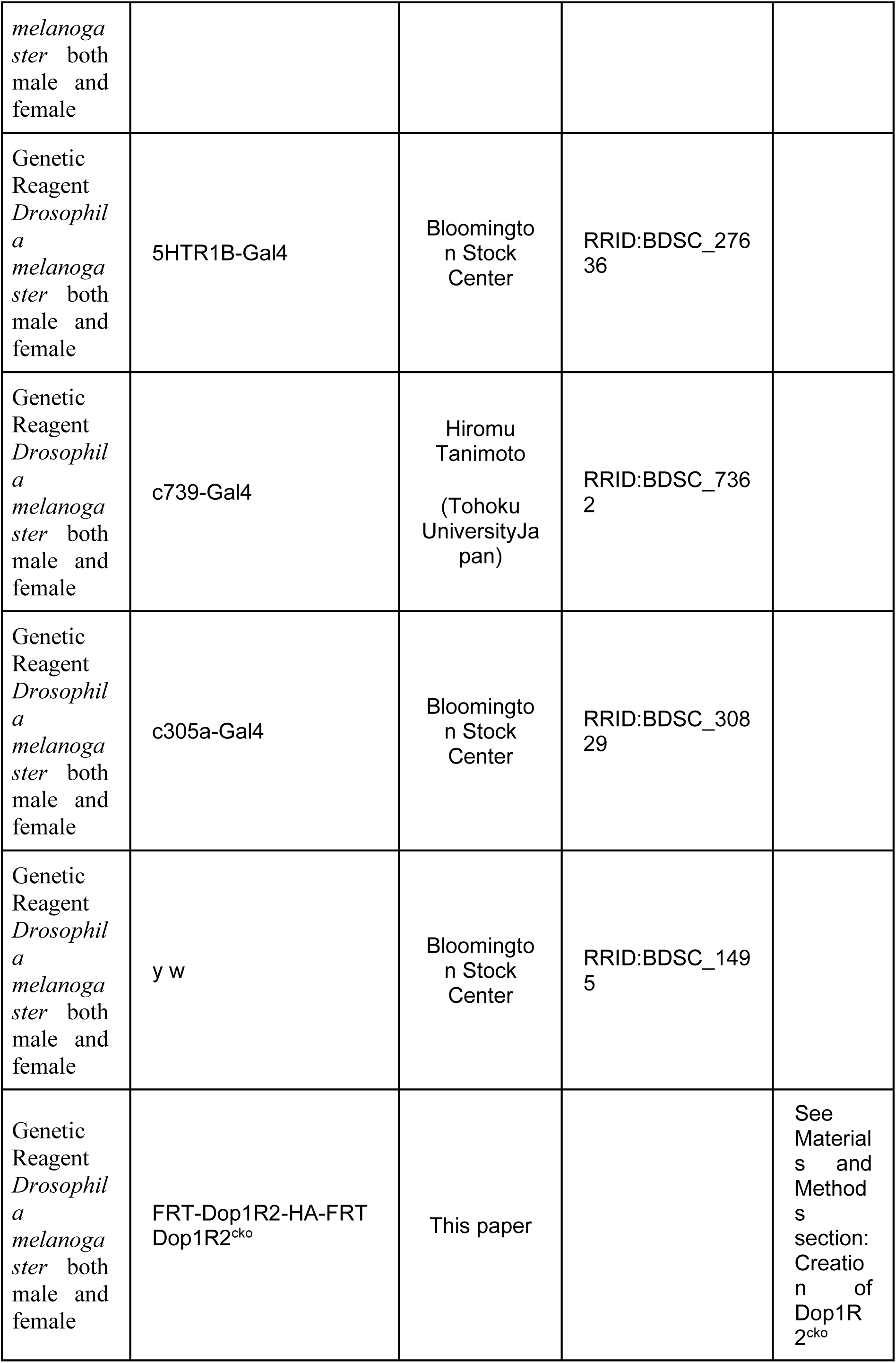
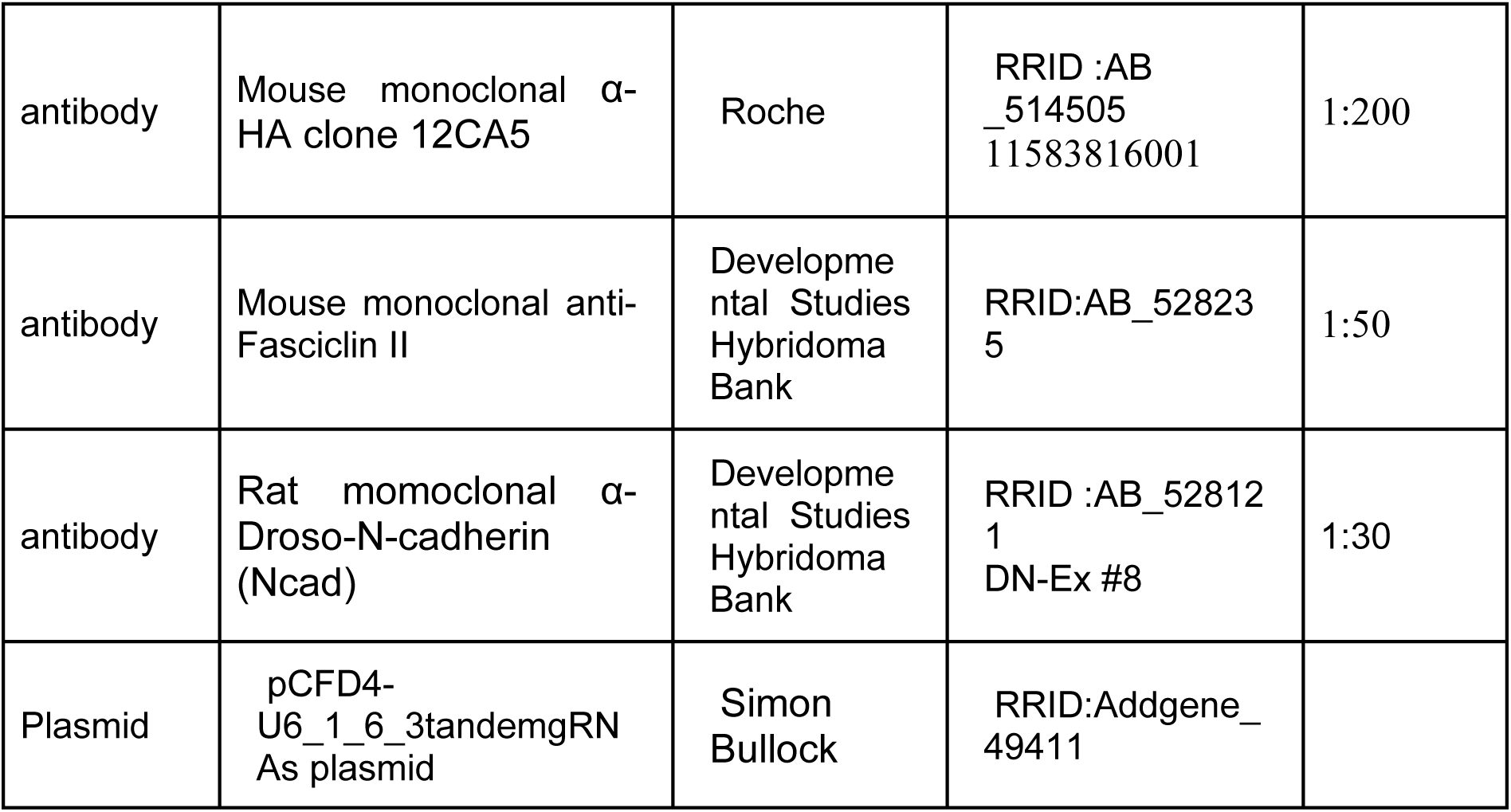
Key Resource Table.

### Fly Husbandry

*Drosophila melanogaster* flies were reared in plastic vials on standard cornmeal food (12g Agar, 40g Sugar, 40g Yeast, 80g Cornmeal per Liter) and transferred to fresh food vials every 2-3 days. Flies were generally kept at 25°C, 60-65% humidity, and exposed to 12 h light and 12 h darkness with light onset at 8 am. The following fly lines were used: *y[1] w[*] P{y[+t7.7]=nos-phiC31\int.NLS}X; P{y[+t7.7]=CaryIP}su(Hw)attP6* (abbreviated *nos>Cas9* in this paper; BL 32232) for microinjection and as PCR template, *w;; Dr e/TM3* (BL 36305) was used as 3rd chromosomal balancer line. *w[1118], 20XUAS-FLPG5.PEST* (BL 55807) was used as *UAS-flp*. OK107*-*Gal4 (106098) was obtained from the Kyoto stock center. The 5HTR1B*-*Gal4 line (BL 27636) and the c305a*-*Gal4 (BL 30829) line are from Bloomington stock center. The c739*-* Gal4 line was gifted to us by H. Tanimoto (Tohoku University Japan). The *y w* (BL 1495) line was used as the control line.

UAS-flp;;Dop1R2 ^cko^ flies and Gal4;Dop1R2 ^cko^ flies were crossed back with ;;Dop ^cko^ flies to obtain appropriate genetic controls which were heterozygous for UAS and Gal4 but not Dop1R2 ^cko^ to control for genetic background.

### Generation of Dop1R2^cko^

For generating the conditional knock-out line we needed three regions: 1) The 5’ flanking sequence upstream of the first FRT site as homologous region, 2) The 3’ flanking sequence downstream of the second FRT site as homologous region, 3) the sequence in between the two FRT sites hereafter named *Dop1R2* coding fragment. To obtain the dopamine receptor fragments, genomic DNA from *nos >Cas9* flies was used as the template for the PCRs. The primers used for the different PCR fragments are shown in Table 2. The 3’ and 5’ fragments of *Dop1R2* were sub-cloned into *pBluescript II SK(+)* vector (*pBS*) with adequate restriction enzymes – SpeI and SmaI for the 5’ fragment and XhoI and Acc65I for the 3’ fragment. The *Dop1R2* coding fragment was cloned with the Invitrogen TOPO kit. After sequence confirmation the fragments were cloned subsequently into *pBS-FRT-3xHA-FRT*. The vector is modified from the vector used by Widmer et al. (Widmer et al., 2018). The GFP was replaced by 3xHA using 3xHA Nco

Fw: (cacatggtTACCCATACGATGTTCCTGACTATGCGGGCTATCCCTATGACGTCCCGGACTATGCAGGATCCTATCCATATGACGTTCCAGATTACGCTgca) and 3xHA H3

Rev: (agcttgcAGCGTAATCTGGAACGTCATATGGATAGGATCCTGCATAGTCCGGGACGTCATAGGGATAGCCCGCATAGTCAGGAACATCGTATGGGTAc) as oligos whereas the FRT site and the pBS backbone were kept. The *Dop1R2* fragment was cloned in-frame in front of the 3xHA tag using AgeI and BstEII-HF as restriction enzymes.

The two guide RNAs were chosen using cripsr optimal target finder (http://targetfinder.flycrispr.neuro.brown.edu/). The used sequences (Table 2) were integrated into PCR primers to amplify a fragment of the *pCFD4-U6_1_6_3tandemgRNAs plasmid* (a gift from Simon Bullock, addgene plasmid # 49411 (Port et al., 2014) that was cloned into BbsI digested *pCFD4-U6_1_6_3tandemgRNAs* vector using Gibbson assembly (NEB). Insertion of the gRNAs was confirmed by sequencing. A mix containing 0.4 mg/ml template and 0.2 mg/ml of the guide RNAs plasmid was injected into *nos>Cas9* embryos. The injected flies were crossed with a 3^rd^ chromosomal balancer line. The F1 generation was crossed again with the 3^rd^ chromosomal balancer flies. As soon as eggs or larvae were visible the adults were sacrificed to check for the *Dop1R2* construct via PCR. Positive hits were then sequenced, and a stock was established.

**Table 2:**
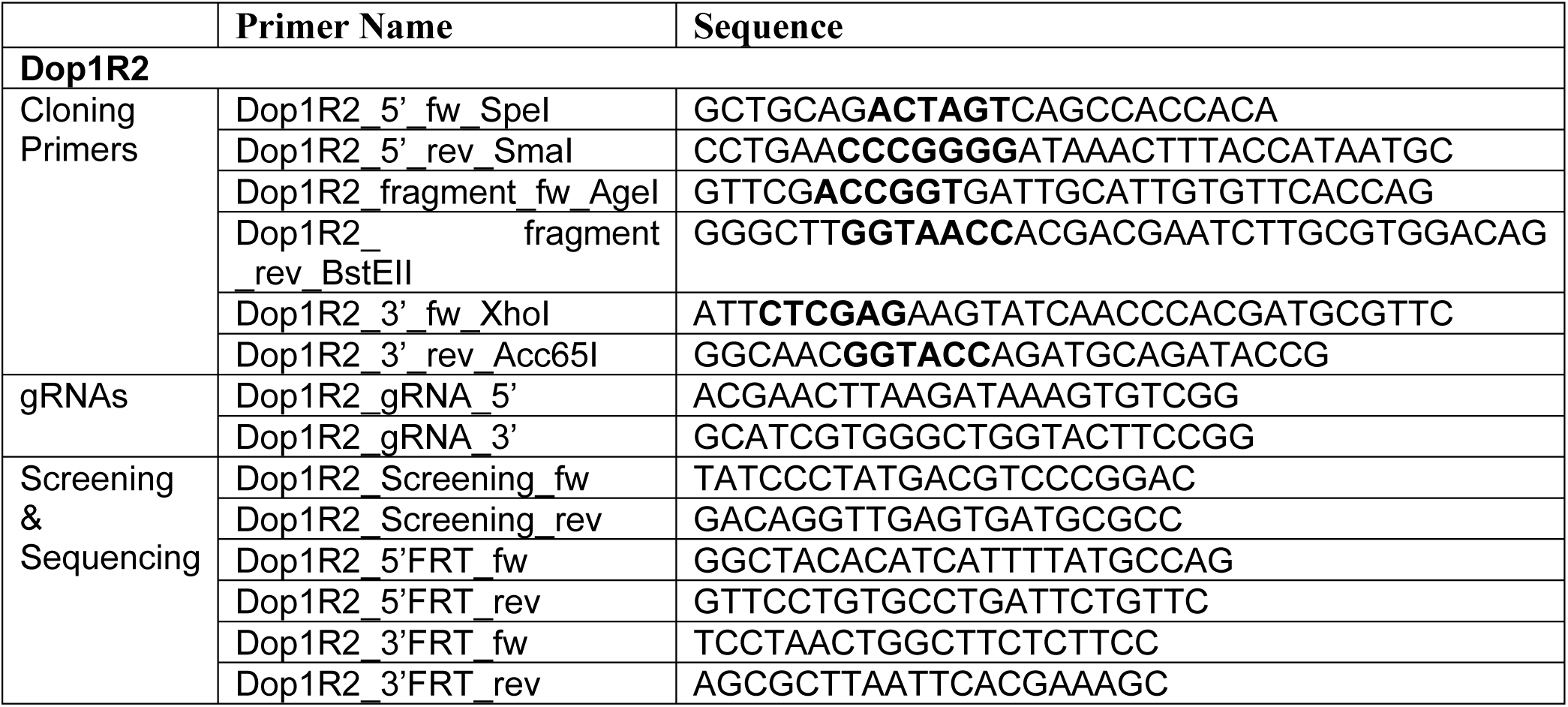
Primers and gRNAs for generating Dop1R2 conditional knock-out lines.

### Immunohistochemistry

Brains from 5–8-day old flies were dissected in PBS (Bio-Froxx 1346LT050) and fixed in 3.7% formaldehyde for 25 min at RT. The brains were washed with 1X PBS containing 0.5% Triton X-100 (Carl Roth 3051.3) (0.5% PBST) before incubating the primary antibodies o/n at 4°C. The primary antibodies were mouse α-HA clone 12CA5 (Roche) 1:200, rat α-Droso-N-cadherin (Ncad) (Iowa H.B: DN-EX8) 1:30 or mouse α-Fasciclin II (FasII) (Iowa H.B: 1D4). After the brains were washed in 0.5% PBST again, they were incubated o/n at 4°C with the secondary antibodies. The following secondary antibodies were used: Goat α-rat Alexa 647 (Molecular probes A-21247) and Goat α-Mouse Alexa 488 (Molecular probes A11029) 1:200. The brains were washed again and mounted in self-made mounting media (90% Glycerol (Fischer Scientific Catalog No. BP229-1), 0.5% N-propyl gallate (Sigma P3130), 20 mM Tris (Fischer Scientific, Catalog No. BP152-5), pH 8.0) (Adapted from NIC Harvard Medical School). The brains were imaged using a confocal microscope (Leica STELLARIS 8 FALCON) at 40X magnification with the Plan APO 40x/1.10 water immersion objective at 1,024 × 1,024 pixels resolution and 600Hz scan rate. Images were processed with Fiji ImageJ and Adobe Illustrator.

### Learning Apparatus

For behavior experiments, we used a memory apparatus that is based on Tully and Quinn’s design and modified to allow conducting four memory experiments in parallel (CON-Elektronik, Greussenheim, Germany). Experiments were performed at 23-25°C and 65-75% relative humidity. The training was performed in dim red light and memory tests were done in complete darkness. The two odors used were 3-octanol (3-Oct) (Sigma-Aldrich 218405) and 4-methyl-cyclohexanol (MCH) (Sigma-Aldrich 66360) diluted in paraffin oil (Sigma-Aldrich 18512) 1:100 respectively. 260μl of the diluted odors were presented in a plastic cup of 14 mm in diameter. A vacuum membrane pump ensured odor delivery at a flow rate of 7 l/min.

### Aversive olfactory conditioning

For aversive conditioning, groups of 50-100 flies of mixed sex were loaded in tubes lined with an electrifiable copper grid. The position in the machine and the sequence in which the genotypes were tested were randomized. Experiments in which more than half of the flies died, the flies did not move or there were technical problems with the machine, as well as human errors were excluded. The training was conducted in the morning. After an accommodation period of 90 s, the first odor was presented for 60 s. In parallel, 12 pulses of 100 V for 1.5 s were delivered with an interval of 3.5 s. After 30 s of flushing with fresh air, the second odor was presented for 60 s. For the subsequent group of flies, the order of the two odors was reversed. For measuring 0 h performance flies were tested about 3 minutes after the end of the conditioning. To determine 2h memory performance, flies were transferred to food vials after conditioning and kept at 25°C until the test. For each genotype and condition the biological replicate is N=12.

For long-term aversive memory, groups of flies were trained with six cycles, where each cycle was composed of 60s of the first odor presented simultaneously with 12 pulses of 90V for 1.5 s with an interval of 3.5s followed by 30s of fresh air and 60s of the second odor. The cycles were spaced by 15 minutes of inter-training-intervals. The flies were starved for 24 hours before testing for long-term memory to increase motivation to climb the Y-Maze and make a decision. The Biological replicate is N=7.

### Appetitive olfactory conditioning

Before appetitive conditioning, groups of 50 to 100 flies with mixed sex were starved for 19 to 21 h in plastic vials containing damp cotton at the bottom. Experiments in which more than half of the flies died, the flies did not move or there were technical problems with the machine, as well as human errors were excluded. The position in the machine and the sequence in which the genotypes were tested were randomized. The training was conducted in the morning. The conditioning protocol consists of a 90-s accommodation period, 120 s of the first odor, 60 s of fresh air followed by 120 s of the second odor. During the first odor, flies are in a conditioning tube lined with filter paper that was soaked in water the day before the experiment and left to dry overnight. For the second odor, flies are transferred to a conditioning tube lined with a filter paper that was soaked with a 1.5 M sucrose (Sigma-Aldrich, Cat# 84100-1KG; CAS Number 57-50-1) solution on the day before and left to dry at RT. After conditioning, flies were either directly tested for STM or put back in starvation vials until the memory test 2 h later. For 24-h memory, flies were fed for 3 h after training before starving them again. One experiment consisted of 2 reciprocal conditionings, in which the odor paired with sucrose was reversed. For each genotype and condition the biological replicate is N=12.

### Memory tests

Flies were loaded into a sliding compartment and transferred to a two-arm choice point. Animals were allowed to choose between 3-octanol and 4-methyl-cyclohexanol. After 60 s, flies trapped in both arms were collected separately and counted. Based on these numbers, a preference index was calculated as follows:

PREF = ((N_arm1_ - N_arm2_) 100) / N_total_ the two preference indices were calculated from the two reciprocal experiments. The average of these two PREFs gives a learning index (LI). LI = (PREF_1_ + PREF_2_) / 2.

In case of all Long-term Aversive memory experiments, the Y-Maze protocol was adapted to test flies 24 hours post training. Testing using the Y-Maze was done following the protocol as described in (Mohandasan et al., 2022) where flies were loaded at the bottom of 20-minutes odorized 3D-printed Y-Mazes from where they would climb up to a choice point and choose between the two odors. The learning index was then calculated after counting the flies in each odorized vial as follows: LI = ((N_CS-_ - N_CS+_) 100) / N_total_. Where N_CS-_ and N_CS+_ are the number of flies that were found trapped in the untrained and trained odor tube respectively.

### Sensory Accuracy tests

Flies were tested for their ability to sense the two used odors 3-octanol and 4-methyl-cyclohexanol as well as electric shock. Therefore, the flies were loaded into a sliding compartment and brought to a two-arm choice point. The flies were allowed to freely choose between an arm containing the stimulus and a neutral arm. All experiments were carried out in the dark. Afterward, the flies in each arm were counted and a preference index was calculated.

For testing the odor response, the flies could choose between one of the odors in the same concentration as used for the behavior experiment and the same amount of paraffin oil for 120s.

Preference index PI = ((N_air_-N_odor_)100)/N_total_.

For shock response, the flies could freely choose for 60s between a cooper-grid lined tube getting pulses of 100V or a cooper-grid lined tube getting no electric shock.

Preference index PI = ((N_No shock_-N_shock_)100)/N_total_.

For testing sugar sensitivity, a group of flies was starved for 1 to 21 h in a tube with damp cotton on the bottom. They could choose for 120 s between a tube lined with filter paper that was soaked in 1.5 M sucrose solution the day before or a tube lined with filter paper that was soaked in distilled water the day before.

Preference index PI = ((N_sucrose_ − N_water_)100) / N_total_

### Statistical analysis

To compare performance indices between different groups we used One-way ANOVA (ANalysis Of VAriance) with posthoc Tukey HSD (Honestly Significant Difference) Test Calculator for comparing multiple treatments in R with the package multcomp. In the case of two groups, we performed a *t*-test for comparison.

To verify if the data was normally distributed, we used Shapiro-Wilk’s test for normal distribution on the statistics software, Prism GraphPad.

We used One-way ANOVA (Analysis of Variance) with posthoc Tukey HSD (Honestly Significant Difference) test when the data was normally distributed. And the non-parametric counterpart, Kruskal-Wallis test when it wasn’t, along with Dunn’s multiple comparisons test. We also verified if all the groups were significantly different from zero using a one sample t-test for normally distributed data and a Wilcoxon’s signed-rank test when the data was not normally distributed.

### Single-cell transcriptomic data analysis

Single-cell Transcriptomics data from (Davie et al., 2018b) was downloaded from the Gene Expression Omnibus (GEO). The metadata containing the annotations was also downloaded and used to annotate the clusters. The clustering and further transcriptomic analyses were performed using the Seurat (version 5.2.1) package for R (Hao et al., 2024). The clustering was done at 0.5 Seurat resolution. And the umap was computed using the top ten principal components. The LogNormalize method from Seurat was used to calculate log-normalized gene expression values. The venn diagram in Supplementary Figure 6 was created using BioVenn (Hulsen et al., 2008).

## References

Akalal D-BG, Yu D, Davis RL. 2011. The Long-Term Memory Trace Formed in the Drosophila α/β Mushroom Body Neurons Is Abolished in Long-Term Memory Mutants. J Neurosci 31:5643–5647. doi:10.1523/JNEUROSCI.3190-10.2011

Alberini CM, Kandel ER. 2015. The Regulation of Transcription in Memory Consolidation. Cold Spring Harb Perspect Biol 7:a021741. doi:10.1101/cshperspect.a021741

Allen AM, Neville MC, Birtles S, Croset V, Treiber CD, Waddell S, Goodwin SF. 2020. A single-cell transcriptomic atlas of the adult Drosophila ventral nerve cord. eLife 9:e54074. doi:10.7554/eLife.54074

Aso Y, Hattori D, Yu Y, Johnston RM, Iyer NA, Ngo T-T, Dionne H, Abbott L, Axel R, Tanimoto H, Rubin GM. 2014a. The neuronal architecture of the mushroom body provides a logic for associative learning. eLife 3:e04577. doi:10.7554/eLife.04577

Aso Y, Herb A, Ogueta M, Siwanowicz I, Templier T, Friedrich AB, Ito K, Scholz H, Tanimoto H. 2012. Three Dopamine Pathways Induce Aversive Odor Memories with Different Stability. PLOS Genetics 8:e1002768. doi:10.1371/journal.pgen.1002768

Aso Y, Sitaraman D, Ichinose T, Kaun KR, Vogt K, Belliart-Guérin G, Plaçais P-Y, Robie AA, Yamagata N, Schnaitmann C, Rowell WJ, Johnston RM, Ngo T-TB, Chen N, Korff W, Nitabach MN, Heberlein U, Preat T, Branson KM, Tanimoto H, Rubin GM. 2014b. Mushroom body output neurons encode valence and guide memory-based action selection in Drosophila. eLife 3:e04580. doi:10.7554/eLife.04580

Berry JA, Cervantes-Sandoval I, Nicholas EP, Davis RL. 2012. Dopamine Is Required for Learning and Forgetting in *Drosophila*. Neuron 74:530–542. doi:10.1016/j.neuron.2012.04.007

Blum AL, Li W, Cressy M, Dubnau J. 2009. Short- and Long-Term Memory in *Drosophila* Require cAMP Signaling in Distinct Neuron Types. Current Biology 19:1341–1350. doi:10.1016/j.cub.2009.07.016

Bouzaiane E, Trannoy S, Scheunemann L, Plaçais P-Y, Preat T. 2015. Two Independent Mushroom Body Output Circuits Retrieve the Six Discrete Components of *Drosophila* Aversive Memory. Cell Reports 11:1280–1292. doi:10.1016/j.celrep.2015.04.044

Burke CJ, Huetteroth W, Owald D, Perisse E, Krashes MJ, Das G, Gohl D, Silies M, Certel S, Waddell S. 2012. Layered reward signalling through octopamine and dopamine in Drosophila. Nature 492:433–437. doi:10.1038/nature11614

Carlezon WA, Duman RS, Nestler EJ. 2005. The many faces of CREB. Trends in Neurosciences 28:436–445. doi:10.1016/j.tins.2005.06.005

Cervantes-Sandoval I, Chakraborty M, MacMullen C, Davis RL. 2016. Scribble Scaffolds a Signalosome for Active Forgetting. Neuron 90:1230–1242. doi:10.1016/j.neuron.2016.05.010

Cervantes-Sandoval I, Martin-Peña A, Berry JA, Davis RL. 2013. System-Like Consolidation of Olfactory Memories in Drosophila. J Neurosci 33:9846–9854. doi:10.1523/JNEUROSCI.0451-13.2013

Cognigni P, Felsenberg J, Waddell S. 2018. Do the right thing: neural network mechanisms of memory formation, expression and update in *Drosophila*. Current Opinion in Neurobiology, Neurobiology of Behavior 49:51–58. doi:10.1016/j.conb.2017.12.002

Crocker A, Guan X-J, Murphy CT, Murthy M. 2016. Cell-Type-Specific Transcriptome Analysis in the *Drosophila* Mushroom Body Reveals Memory-Related Changes in Gene Expression. Cell Reports 15:1580–1596. doi:10.1016/j.celrep.2016.04.046

Croset V, Treiber CD, Waddell S. 2018. Cellular diversity in the Drosophila midbrain revealed by single-cell transcriptomics. eLife 7:e34550. doi:10.7554/eLife.34550

Davie K, Janssens J, Koldere D, De Waegeneer M, Pech U, Kreft Ł, Aibar S, Makhzami S, Christiaens V, Bravo González-Blas C, Poovathingal S, Hulselmans G, Spanier KI, Moerman T, Vanspauwen B, Geurs S, Voet T, Lammertyn J, Thienpont B, Liu S, Konstantinides N, Fiers M, Verstreken P, Aerts S. 2018. A Single-Cell Transcriptome Atlas of the Aging Drosophila Brain. Cell 174:982–998.e20. doi:10.1016/j.cell.2018.05.057

de Queiroz BR, Laghrissi H, Rajeev S, Blot L, De Graeve F, Dehecq M, Hallegger M, Dag U, Dunoyer de Segonzac M, Ramialison M, Cazevieille C, Keleman K, Ule J, Hubstenberger A, Besse F. 2025. Axonal RNA localization is essential for long-term memory. Nat Commun 16:2560. doi:10.1038/s41467-025-57651-7

Feng G, Hannan F, Reale V, Hon YY, Kousky CT, Evans PD, Hall LM. 1996. Cloning and Functional Characterization of a Novel Dopamine Receptor from Drosophila melanogaster. J Neurosci 16:3925–3933. doi:10.1523/JNEUROSCI.16-12-03925.1996

Ferris J, Ge H, Liu L, Roman G. 2006. G(o) signaling is required for Drosophila associative learning. Nat Neurosci 9:1036–1040. doi:10.1038/nn1738

Golic KG, Lindquist S. 1989. The FLP recombinase of yeast catalyzes site-specific recombination in the drosophila genome. Cell 59:499–509. doi:10.1016/0092-8674(89)90033-0

Gratz SJ, Ukken FP, Rubinstein CD, Thiede G, Donohue LK, Cummings AM, O’Connor-Giles KM. 2014. Highly Specific and Efficient CRISPR/Cas9-Catalyzed Homology-Directed Repair in Drosophila. Genetics 196:961–971. doi:10.1534/genetics.113.160713

Guven-Ozkan T, Davis RL. 2014. Functional neuroanatomy of Drosophila olfactory memory formation. Learn Mem 21:519–526. doi:10.1101/lm.034363.114

Han K-A, Millar NS, Grotewiel MS, Davis RL. 1996. DAMB, a Novel Dopamine Receptor Expressed Specifically in Drosophila Mushroom Bodies. Neuron 16:1127–1135. doi:10.1016/S0896-6273(00)80139-7

Handler A, Graham TGW, Cohn R, Morantte I, Siliciano AF, Zeng J, Li Y, Ruta V. 2019. Distinct Dopamine Receptor Pathways Underlie the Temporal Sensitivity of Associative Learning. Cell 178:60–75.e19. doi:10.1016/j.cell.2019.05.040

Hao Y, Stuart T, Kowalski MH, Choudhary S, Hoffman P, Hartman A, Srivastava A, Molla G, Madad S, Fernandez-Granda C, Satija R. 2024. Dictionary learning for integrative, multimodal and scalable single-cell analysis. Nat Biotechnol 42:293–304. doi:10.1038/s41587-023-01767-y

Hearn MG, Ren Y, McBride EW, Reveillaud I, Beinborn M, Kopin AS. 2002. A Drosophila dopamine 2-like receptor: Molecular characterization and identification of multiple alternatively spliced variants. Proceedings of the National Academy of Sciences 99:14554–14559. doi:10.1073/pnas.202498299

Hermans E. 2003. Biochemical and pharmacological control of the multiplicity of coupling at G-protein-coupled receptors. Pharmacol Ther 99:25–44. doi:10.1016/s0163-7258(03)00051-2

Himmelreich S, Masuho I, Berry JA, MacMullen C, Skamangas NK, Martemyanov KA, Davis RL. 2017. Dopamine Receptor DAMB Signals via Gq to Mediate Forgetting in *Drosophila*. Cell Reports 21:2074–2081. doi:10.1016/j.celrep.2017.10.108

Huang C, Zheng X, Zhao H, Li M, Wang P, Xie Z, Wang L, Zhong Y. 2012. A Permissive Role of Mushroom Body α/β Core Neurons in Long-Term Memory Consolidation in *Drosophila*. Current Biology 22:1981–1989. doi:10.1016/j.cub.2012.08.048

Hulsen T, de Vlieg J, Alkema W. 2008. BioVenn – a web application for the comparison and visualization of biological lists using area-proportional Venn diagrams. BMC Genomics 9:488. doi:10.1186/1471-2164-9-488

Ichinose T, Aso Y, Yamagata N, Abe A, Rubin GM, Tanimoto H. 2015. Reward signal in a recurrent circuit drives appetitive long-term memory formation. eLife 4:e10719. doi:10.7554/eLife.10719

Iversen SD, Iversen LL. 2007. Dopamine: 50 years in perspective. Trends Neurosci 30:188–193. doi:10.1016/j.tins.2007.03.002

Joga MR, Zotti MJ, Smagghe G, Christiaens O. 2016. RNAi Efficiency, Systemic Properties, and Novel Delivery Methods for Pest Insect Control: What We Know So Far. Front Physiol 7. doi:10.3389/fphys.2016.00553

Kaldun JC, Sprecher SG. 2019. Initiated by CREB: Resolving Gene Regulatory Programs in Learning and Memory. BioEssays 41:1900045. doi:10.1002/bies.201900045

Kandel ER. 2012. The molecular biology of memory: cAMP, PKA, CRE, CREB-1, CREB-2, and CPEB. Molecular Brain 5:14. doi:10.1186/1756-6606-5-14

Karam CS, Jones SK, Javitch JA. 2020. Come Fly with Me: An overview of dopamine receptors in Drosophila melanogaster. Basic & Clinical Pharmacology & Toxicology 126:56–65. doi:10.1111/bcpt.13277

Kim Y-C, Lee H-G, Han K-A. 2007. D1 Dopamine Receptor dDA1 Is Required in the Mushroom Body Neurons for Aversive and Appetitive Learning in Drosophila. J Neurosci 27:7640–7647. doi:10.1523/JNEUROSCI.1167-07.2007

Kim Y-C, Lee H-G, Seong C-S, Han K-A. 2003. Expression of a D1 dopamine receptor dDA1/DmDOP1 in the central nervous system of *Drosophila melanogaster*. Gene Expression Patterns 3:237–245. doi:10.1016/S1567-133X(02)00098-4

Klein MO, Battagello DS, Cardoso AR, Hauser DN, Bittencourt JC, Correa RG. 2019. Dopamine: Functions, Signaling, and Association with Neurological Diseases. Cell Mol Neurobiol 39:31–59. doi:10.1007/s10571-018-0632-3

Krashes MJ, Waddell S. 2008. Rapid Consolidation to a radish and Protein Synthesis-Dependent Long-Term Memory after Single-Session Appetitive Olfactory Conditioning in Drosophila. J Neurosci 28:3103–3113. doi:10.1523/JNEUROSCI.5333-07.2008

Lark A, Kitamoto T, Martin J-R. 2017. Modulation of neuronal activity in the *Drosophila* mushroom body by DopEcR, a unique dual receptor for ecdysone and dopamine. Biochimica et Biophysica Acta (BBA) - Molecular Cell Research 1864:1578–1588. doi:10.1016/j.bbamcr.2017.05.015

Ledonne A, Mercuri NB. 2017. Current Concepts on the Physiopathological Relevance of Dopaminergic Receptors. Front Cell Neurosci 11. doi:10.3389/fncel.2017.00027

Missale C, Nash SR, Robinson SW, Jaber M, Caron MG. 1998. Dopamine Receptors: From Structure to Function. Physiological Reviews 78:189–225. doi:10.1152/physrev.1998.78.1.189

Miyashita T, Kikuchi E, Horiuchi J, Saitoe M. 2018. Long-Term Memory Engram Cells Are Established by c-Fos/CREB Transcriptional Cycling. Cell Reports 25:2716–2728.e3. doi:10.1016/j.celrep.2018.11.022

Mohandasan R, Thakare M, Sunke S, Iqbal FM, Sridharan M, Das G. 2022. Enhanced olfactory memory detection in trap-design Y-mazes allows the study of imperceptible memory traces in Drosophila. Learn Mem 29:355–366. doi:10.1101/lm.053545.121

Musso P-Y, Tchenio P, Preat T. 2015. Delayed Dopamine Signaling of Energy Level Builds Appetitive Long-Term Memory in *Drosophila*. Cell Reports 10:1023–1031. doi:10.1016/j.celrep.2015.01.036

Neves SR, Ram PT, Iyengar R. 2002. G Protein Pathways. Science 296:1636–1639. doi:10.1126/science.1071550

Nishi A, Kuroiwa M, Shuto T. 2011. Mechanisms for the Modulation of Dopamine D1 Receptor Signaling in Striatal Neurons. Front Neuroanat 5. doi:10.3389/fnana.2011.00043

Owald D, Felsenberg J, Talbot CB, Das G, Perisse E, Huetteroth W, Waddell S. 2015. Activity of Defined Mushroom Body Output Neurons Underlies Learned Olfactory Behavior in *Drosophila*. Neuron 86:417–427. doi:10.1016/j.neuron.2015.03.025

Pavlowsky A, Schor J, Plaçais P-Y, Preat T. 2018. A GABAergic Feedback Shapes Dopaminergic Input on the *Drosophila* Mushroom Body to Promote Appetitive Long-Term Memory. Current Biology 28:1783–1793.e4. doi:10.1016/j.cub.2018.04.040

Perisse E, Burke C, Huetteroth W, Waddell S. 2013. Shocking Revelations and Saccharin Sweetness in the Study of *Drosophila* Olfactory Memory. Current Biology 23:R752–R763. doi:10.1016/j.cub.2013.07.060

Plaçais P-Y, de Tredern É, Scheunemann L, Trannoy S, Goguel V, Han K-A, Isabel G, Preat T. 2017. Upregulated energy metabolism in the Drosophila mushroom body is the trigger for long-term memory. Nat Commun 8:15510. doi:10.1038/ncomms15510

Plaçais P-Y, Preat T. 2013. To Favor Survival Under Food Shortage, the Brain Disables Costly Memory. Science 339:440–442. doi:10.1126/science.1226018

Plaçais P-Y, Trannoy S, Friedrich AB, Tanimoto H, Preat T. 2013. Two pairs of mushroom body efferent neurons are required for appetitive long-term memory retrieval in Drosophila. Cell Rep 5:769–780. doi:10.1016/j.celrep.2013.09.032

Plaçais P-Y, Trannoy S, Isabel G, Aso Y, Siwanowicz I, Belliart-Guérin G, Vernier P, Birman S, Tanimoto H, Preat T. 2012. Slow oscillations in two pairs of dopaminergic neurons gate long-term memory formation in Drosophila. Nat Neurosci 15:592–599. doi:10.1038/nn.3055

Port F, Chen H-M, Lee T, Bullock SL. 2014. Optimized CRISPR/Cas tools for efficient germline and somatic genome engineering in Drosophila. Proceedings of the National Academy of Sciences 111:E2967–E2976. doi:10.1073/pnas.1405500111

Qi C, Lee D. 2014. Pre- and Postsynaptic Role of Dopamine D2 Receptor DD2R in Drosophila Olfactory Associative Learning. Biology 3:831–845. doi:10.3390/biology3040831

Qin H, Cressy M, Li W, Coravos JS, Izzi SA, Dubnau J. 2012. Gamma Neurons Mediate Dopaminergic Input during Aversive Olfactory Memory Formation in *Drosophila*. Current Biology 22:608–614. doi:10.1016/j.cub.2012.02.014

Raun N, Jones SG, Kerr O, Keung C, Butler EF, Alka K, Krupski JD, Reid-Taylor RA, Ibrahim V, Williams M, Top D, Kramer JM. 2025. Trithorax regulates long-term memory in Drosophila through epigenetic maintenance of mushroom body metabolic state and translation capacity. PLOS Biology 23:e3003004. doi:10.1371/journal.pbio.3003004

Sabandal JM, Berry JA, Davis RL. 2021. Dopamine-based mechanism for transient forgetting. Nature 591:426–430. doi:10.1038/s41586-020-03154-y

Sabandal JM, Sabandal PR, Kim Y-C, Han K-A. 2020. Concerted Actions of Octopamine and Dopamine Receptors Drive Olfactory Learning. J Neurosci 40:4240–4250. doi:10.1523/JNEUROSCI.1756-19.2020

Scheunemann L, Jost E, Richlitzki A, Day JP, Sebastian S, Thum AS, Efetova M, Davies S-A, Schwärzel M. 2012. Consolidated and Labile Odor Memory Are Separately Encoded within the Drosophila Brain. J Neurosci 32:17163–17171. doi:10.1523/JNEUROSCI.3286-12.2012

Scholz-Kornehl S, Schwärzel M. 2016. Circuit Analysis of a Drosophila Dopamine Type 2 Receptor That Supports Anesthesia-Resistant Memory. J Neurosci 36:7936–7945. doi:10.1523/JNEUROSCI.4475-15.2016

Shuai Y, Lu B, Hu Y, Wang L, Sun K, Zhong Y. 2010. Forgetting Is Regulated through Rac Activity in *Drosophila*. Cell 140:579–589. doi:10.1016/j.cell.2009.12.044

Shyu W-H, Lee W-P, Chiang M-H, Chang C-C, Fu T-F, Chiang H-C, Wu T, Wu C-L. 2019. Electrical synapses between mushroom body neurons are critical for consolidated memory retrieval in Drosophila. PLOS Genetics 15:e1008153. doi:10.1371/journal.pgen.1008153

Siju KP, De Backer J-F, Grunwald Kadow IC. 2021. Dopamine modulation of sensory processing and adaptive behavior in flies. Cell Tissue Res 383:207–225. doi:10.1007/s00441-020-03371-x

Siju KP, Štih V, Aimon S, Gjorgjieva J, Portugues R, Grunwald Kadow IC. 2020. Valence and State-Dependent Population Coding in Dopaminergic Neurons in the Fly Mushroom Body. Current Biology 30:2104–2115.e4. doi:10.1016/j.cub.2020.04.037

Sitaraman D, Aso Y, Rubin GM, Nitabach MN. 2015. Control of Sleep by Dopaminergic Inputs to the Drosophila Mushroom Body. Front Neural Circuits 9. doi:10.3389/fncir.2015.00073

Srivastava DP, Yu EJ, Kennedy K, Chatwin H, Reale V, Hamon M, Smith T, Evans PD. 2005. Rapid, Nongenomic Responses to Ecdysteroids and Catecholamines Mediated by a Novel Drosophila G-Protein-Coupled Receptor. J Neurosci 25:6145–6155. doi:10.1523/JNEUROSCI.1005-05.2005

Sugamori KS, Demchyshyn LL, McConkey F, Forte MA, Niznik HB. 1995. A primordial dopamine D1-like adenylyl cyclase-linked receptor from Drosophila melanogaster displaying poor affinity for benzazepines. FEBS Lett 362:131–138. doi:10.1016/0014-5793(95)00224-w

Sun H, Nishioka T, Hiramatsu S, Kondo S, Amano M, Kaibuchi K, Ichinose T, Tanimoto H. 2020. Dopamine Receptor Dop1R2 Stabilizes Appetitive Olfactory Memory through the Raf/MAPK Pathway in Drosophila. J Neurosci 40:2935–2942. doi:10.1523/JNEUROSCI.1572-19.2020

Tanaka NK, Tanimoto H, Ito K. 2008. Neuronal assemblies of the Drosophila mushroom body. Journal of Comparative Neurology 508:711–755. doi:10.1002/cne.21692

Tomita J, Ban G, Kume K. 2017. Genes and neural circuits for sleep of the fruit fly. Neuroscience Research, Cutting-edge Approaches to Unwrapping the Mysteries of Sleep 118:82–91. doi:10.1016/j.neures.2017.04.010

Trannoy S, Redt-Clouet C, Dura J-M, Preat T. 2011. Parallel Processing of Appetitive Short- and Long-Term Memories In *Drosophila*. Current Biology 21:1647–1653. doi:10.1016/j.cub.2011.08.032

Tritsch NX, Sabatini BL. 2012. Dopaminergic Modulation of Synaptic Transmission in Cortex and Striatum. Neuron 76:33–50. doi:10.1016/j.neuron.2012.09.023

Tully T, Preat T, Boynton SC, Del Vecchio M. 1994. Genetic dissection of consolidated memory in Drosophila. Cell 79:35–47. doi:10.1016/0092-8674(94)90398-0

Turrel O, Ramesh N, Escher MJF, Pooryasin A, Sigrist SJ. 2022. Transient active zone remodeling in the *Drosophila* mushroom body supports memory. Current Biology 32:4900–4913.e4. doi:10.1016/j.cub.2022.10.017

Ueoka Y, Hiroi M, Abe T, Tabata T. 2017. Suppression of a single pair of mushroom body output neurons in Drosophila triggers aversive associations. FEBS Open Bio 7:562–576. doi:10.1002/2211-5463.12203

Waddell S. 2013. Reinforcement signalling in *Drosophila*; dopamine does it all after all. Current Opinion in Neurobiology, Social and emotional neuroscience 23:324–329. doi:10.1016/j.conb.2013.01.005

Widmer YF, Fritsch C, Jungo MM, Almeida S, Egger B, Sprecher SG. 2018. Multiple neurons encode CrebB dependent appetitive long-term memory in the mushroom body circuit. eLife 7:e39196. doi:10.7554/eLife.39196

Yamamoto S, Seto ES. 2014. Dopamine Dynamics and Signaling in Drosophila: An Overview of Genes, Drugs and Behavioral Paradigms. Experimental Animals 63:107–119. doi:10.1538/expanim.63.107

Yang Q, Zhou J, Wang L, Hu W, Zhong Y, Li Q. 2023. Spontaneous recovery of reward memory through active forgetting of extinction memory. Current Biology 33:838–848.e3. doi:10.1016/j.cub.2023.01.022

Zars T, Fischer † M., Schulz R, Heisenberg M. 2000. Localization of a Short-Term Memory in Drosophila. Science 288:672–675. doi:10.1126/science.288.5466.672

Zhou M, Chen N, Tian J, Zeng J, Zhang Y, Zhang X, Guo J, Sun J, Li Yulong, Guo A, Li Yan. 2019. Suppression of GABAergic neurons through D2-like receptor secures efficient conditioning in Drosophila aversive olfactory learning. Proceedings of the National Academy of Sciences 116:5118–5125. doi:10.1073/pnas.1812342116

